# A chromosomal inversion drives evolution of multiple adaptive traits in deer mice

**DOI:** 10.1101/2021.01.21.427490

**Authors:** Emily R. Hager, Olivia S. Harringmeyer, T. Brock Wooldridge, Shunn Theingi, Jacob T. Gable, Sade McFadden, Beverly Neugeboren, Kyle M. Turner, Hopi E. Hoekstra

## Abstract

A long-standing question in evolutionary biology is how differences in multiple traits can evolve quickly and be maintained together during local adaptation. Using forest and prairie ecotypes in deer mice, which differ in both tail length and coat color, we discovered a 41 Mb chromosomal inversion that is strongly linked to variation in both traits. The inversion maintains highly divergent loci in strong linkage disequilibrium and likely originated ~170 kya, long before the forest-prairie divergence ~10 kya. Consistent with a role in local adaptation, inversion frequency is associated with phenotype and habitat across both a local transect and the species range. Still, although eastern and western forest subspecies share similar phenotypes, the inversion is absent in eastern North America. This work highlights the significance of inversion polymorphisms for the establishment and maintenance of multiple locally adaptive traits in mammals, and demonstrates that, even within a species, parallel phenotypes may evolve through nonparallel genetic mechanisms.

---

Rapid adaptation to novel environments often involves concurrent divergence in multiple traits. Determining how these trait differences are generated and maintained together in the face of gene flow is critical to understanding local adaptation. To address this goal, we focused on two distinct ecotypes, first described by natural historians in the early 1900’s, that occur within a single species, the North American deer mouse *(Peromyscus maniculatus):* a semi-arboreal forest form and a terrestrial prairie form that differ in multiple traits, including tail length and coat color (*1–3*). Longer tails are thought to have evolved as an adaptation to an arboreal lifestyle in forested habitat (*1, 2, 4, 5*), while coat color in deer mice often evolves to match the local soil substrate, likely through predation-mediated selection (*2, 6–8*). The forest ecotype is thought to have evolved multiple times since the last glacial maximum ~12 thousand years ago (kya, (*9, 10*)) and is maintained despite ongoing gene flow from neighboring prairie populations (*11*).

## Forest and prairie mice differ in multiple traits

We selected focal populations from coastal temperate rainforest (*P. m. rubidus,* ‘forest’ ecotype) and arid sagebrush steppe (*P. m. gambelii,* ‘prairie’ ecotype) habitat, separated by approximately 500 km (Figure 1A; (*11, 12*)). We established laboratory colonies from wild-caught mice and took standard body measurements (tail, body, hindfoot and ear lengths as well as weight) and coat color measurements (brightness, hue and saturation in three body regions) in both wild-caught mice and their laboratory-reared descendants. Forest mice consistently had longer tails, longer hind feet, and darker, redder coats than prairie mice (Figure 1B,C, Figure S1, Table S1), and these differences persisted in laboratory-born mice raised in common conditions (Figure S2, Table S1), suggesting a strong genetic component to these classic forest phenotypes.

**Fig. 1.**
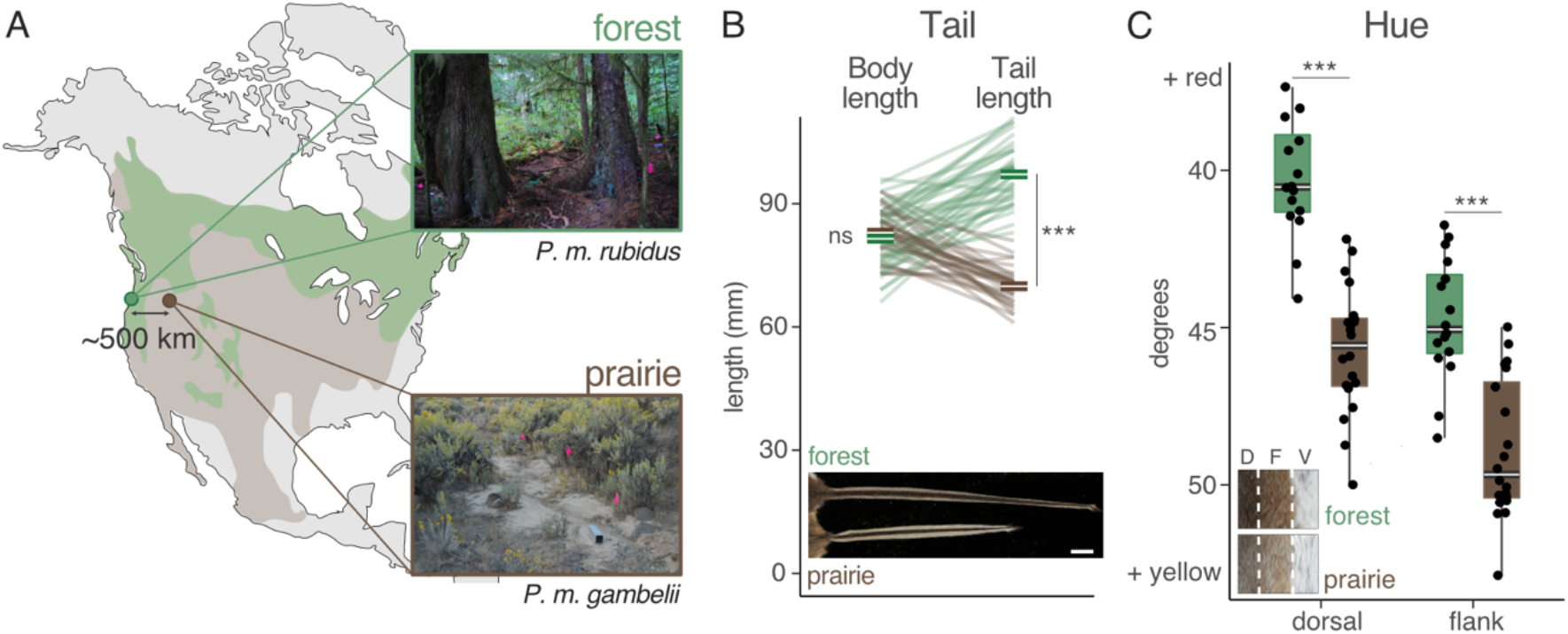
Forest and prairie mice differ in tail length and pigmentation. (A) We measured wild-caught forest (*P. m. rubidus,* green) and prairie (*P. m. gambelii,* brown) ecotypes from western and eastern Oregon, USA, respectively. Map shows the approximate range of forest (green) and prairie (brown) ecotypes in North America. Photos show representative capture sites; pink flags indicate trap lines. (B) Body length (left; not including the tail) and tail length (right) for wild-caught adult mice (n = 38 forest, 32 prairie). Lines connect body and tail measurements for the same individual. Inset: image of a representative tail from each ecotype (scale bar = 1 cm). (C) Hue values for the dorsal and flank regions of wild-caught adult mice (n = 16 forest, 20 prairie). Inset: Dorsal (D), flank (F), and ventral (V) regions from a representative forest and prairie mouse. Photos in (B,C): Museum of Comparative Zoology, Harvard University. Symbols: ns=p>0.05; ***=p<0.001 (Welch’s t-test, two-sided).

## A large inversion is associated with tail length and coat color

Using a forward-genetic approach, we identified genomic regions linked to these ecotype-specific differences in morphology. Specifically, we intercrossed forest and prairie mice in the lab to generate 555 second-generation (F2) hybrids (forest female x prairie male, n = 203 F2s, and prairie female x forest male, n = 352 F2s) and inferred hybrid genotypes (*13*) with sequence data generated by double-digest restriction associated DNA sequencing (*14*) before performing quantitative trait locus (QTL) mapping for each trait (Figure 2, Figure S3, Table S2). We identified five regions associated with tail length differences; at each locus, forest ancestry was associated with longer tails (total percent variance explained (PVE): 27%; individual PVE: 2.6 – 12.1%). Only one region, on chromosome 15, was strongly and significantly associated with coat color (PVE, dorsal hue: 40.0%; flank hue: 45.6%), and it overlapped with the largest-effect locus associated with tail length (95% Bayes credible interval: dorsal hue = 0.4-40.5 Mb; flank hue = 0.4-39.4 Mb; tail length = 0.4-41.5 Mb). Thus, a single region on chromosome 15 was strongly associated with ecotype-specific differences in both tail length and coat color.

**Fig. 2.**
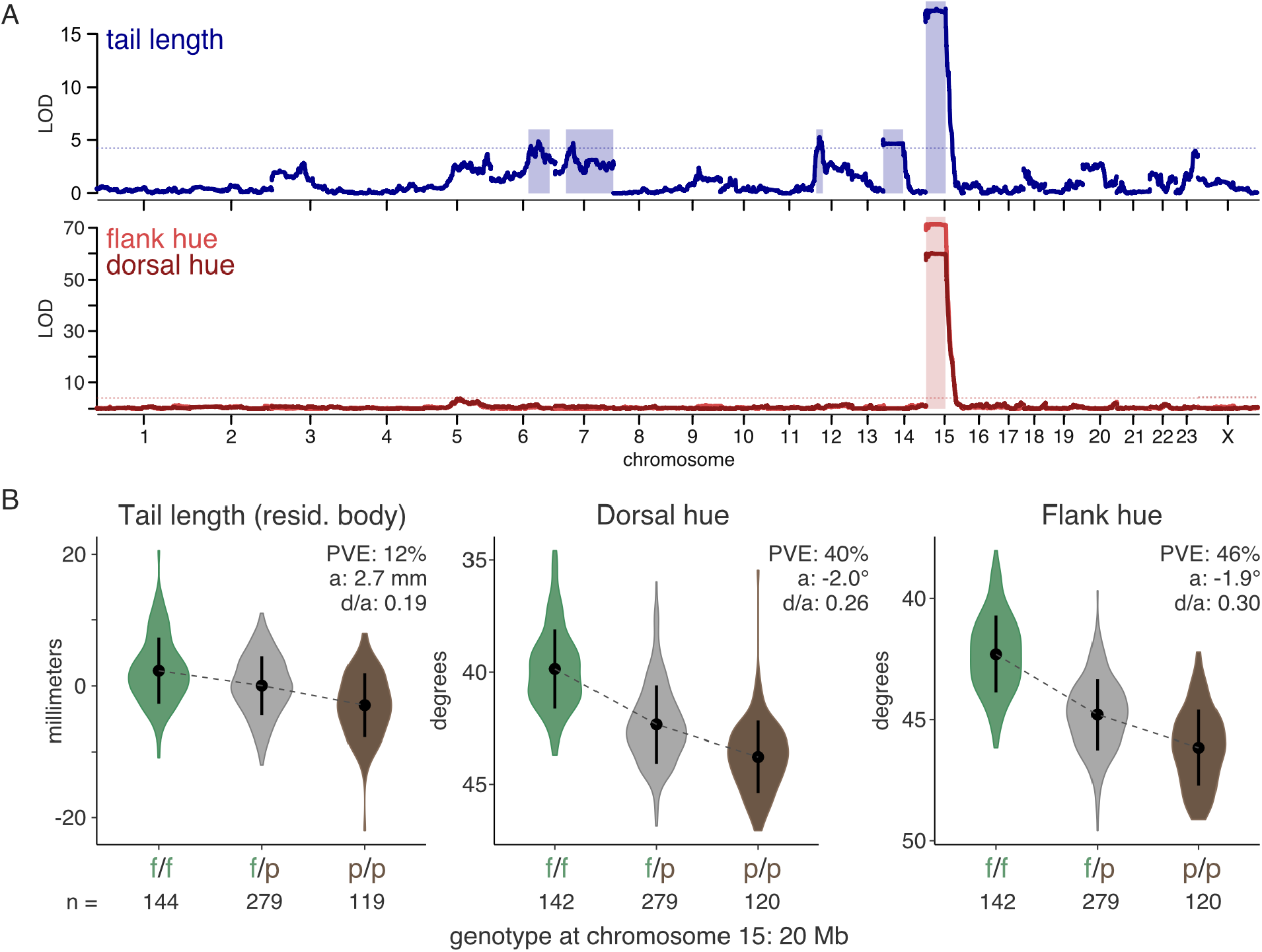
A region on chromosome 15 is strongly associated with both tail length and hue. (A) Statistical association (LOD, or log of the odds, score) of ancestry with tail length (top, blue) and dorsal and flank hue (bottom; dorsal = dark red, flank = light red) in laboratory-reared F2 hybrids (tail, n = 542; hue, n = 541). Physical distance (in basepairs) is shown on the x-axis; axis labels indicate the center of each chromosome. Dotted lines indicate the genome-wide significance threshold (a = 0.05) based on permutation tests, and shaded rectangles indicate the 95% Bayes’ credible intervals for all chromosomes with significant QTL peaks. For tail length analysis, body length was included as an additive covariate. (B) Tail length (left, shown after taking the residual against body length in the hybrids), dorsal hue (center) and flank hue (right) of F2 hybrids, binned by genotype at 20 Mb on chromosome 15 (f/f = homozygous forest; f/p = heterozygous; p/p = homozygous prairie; sample sizes as shown in figure). Points and error bars show mean ± standard deviation. PVE = percent of the variance explained by genotype. a = additive effect of one forest allele. d/a = absolute value of the dominance ratio.

The QTL peak on chromosome 15 exhibited an unusual pattern of association with both morphological traits. Specifically, the strength of association between genotype and phenotype remained largely constant across half the chromosome (Figure 3A). This pattern reflects reduced recombination between forest and prairie alleles in the laboratory cross: only 2 of 1110 F2 chromosomes were recombinant in this region (Figure 3B). In addition, we found consistently elevated F_ST_ (Figure 3C) and high linkage disequilibrium (Figure 3D) across this genetic region in wild populations (whole genome re-sequencing: n = 15 forest, 15 prairie). Together, these data are consistent with little to no recombination across half of chromosome 15 both in the laboratory and wild populations.

**Fig. 3.**
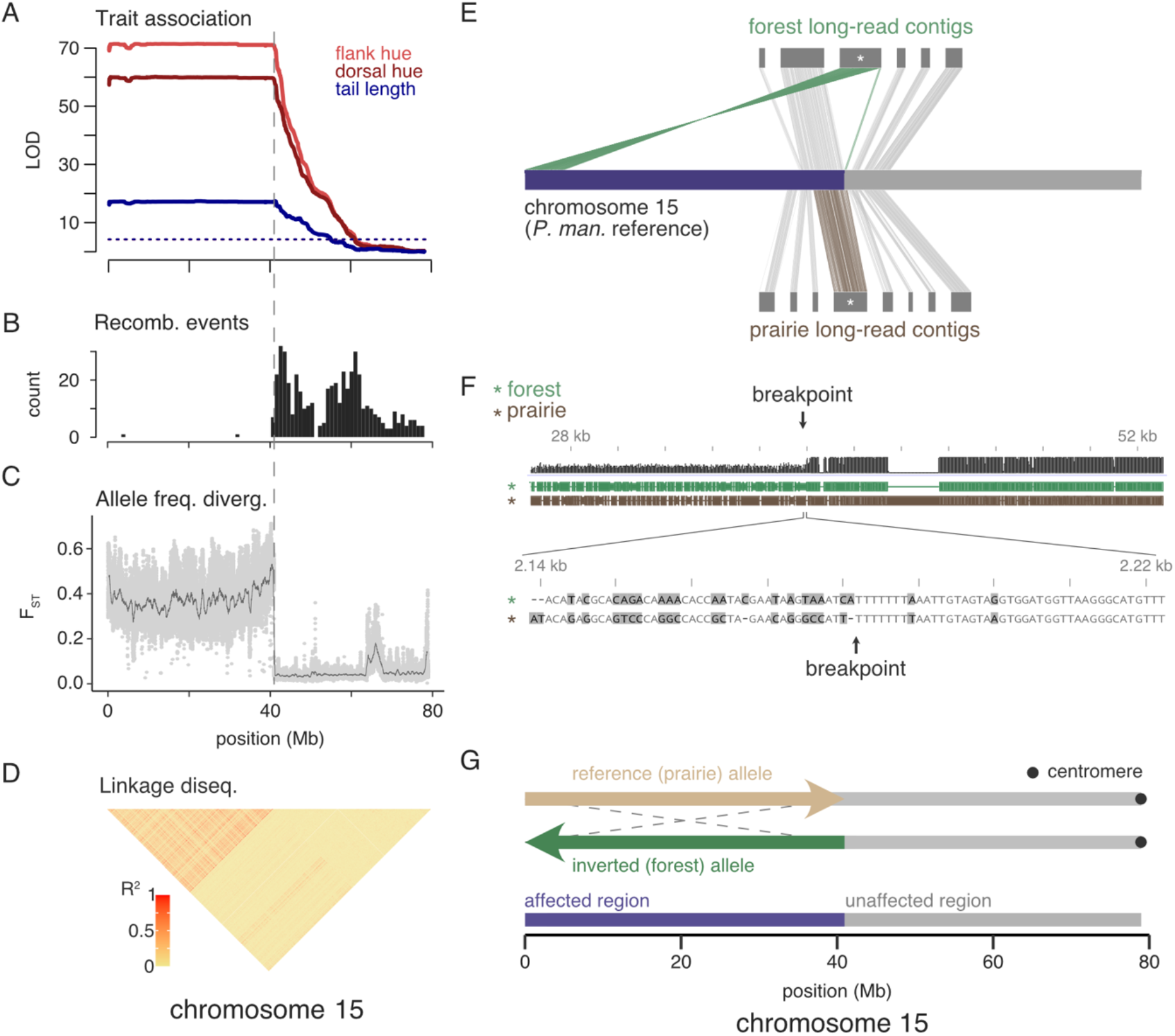
The region associated with tail length and hue is a large chromosomal inversion. Across chromosome 15: (A-B) show data in F2 hybrids; (C-D) show data in wild-caught mice (n = 15 forest, 15 prairie). (A) LOD score for tail length (blue), dorsal hue (dark red) and flank hue (light red). (B) Number of recombination breakpoint events, binned in 1-Mb windows. (C) *F_ST_* between forest and prairie mice estimated in 10-kb windows with step size of 1 kb (light gray dots). Dark gray line shows data smoothed with a moving average over 500 windows. (D) Linkage disequilibrium across forest and prairie mice. Heatmap shows r^2^ computed between genotypes at SNPs with minor allele frequency greater than 0.1 and thinned to 1 SNP per 100 kb. (E) Contigs assembled from PacBio long-read sequencing for one forest (top) and one prairie (bottom) mouse. Only contigs that fully or partially mapped to the *P. maniculatus* reference genome from chr15:35-45 Mb and were > 0.5 Mb in length are shown. Starred contigs localize the inversion breakpoint (chr15: 40.94 Mb), with a single prairie contig (brown) mapping continuously from chr15: 37.0-41.3 Mb whereas a single forest contig (green) maps continuously from chr15: 5.1-0 Mb then chr15: 40.94-41.2 Mb. The region of chromosome 15 affected by the inversion is highlighted in purple. (F) Alignment between regions of the forest and prairie contigs surrounding the breakpoint (top: black = alignment quality, green = forest contig, brown = prairie contig). Large prairie insertion near the breakpoint is a transposon. Bottom: basepair-level alignment around the breakpoint; gray = mismatch. (G) Model of the inverted (green) and reference (tan) alleles. The inversion spans 0-40.9 Mb (affected region, purple), and excludes 40.9-79 Mb (unaffected region, gray). Mapping common centromere-like sequence repeats to both sets of contigs localized the likely centromere to the end of the unaffected region.

This pattern of reduced recombination could be produced by a large genomic rearrangement (or a set of rearrangements). To determine the nature of any rearrangements on chromosome 15, we used PacBio long-read sequencing (n = 1 forest, 1 prairie) (*15*). First, we generated independent *de novo* assemblies for each individual and mapped the resulting contigs to the reference genome for *P. m. bairdii.* In the forest individual, one contig mapped near the center of the chromosome (from 41.19-40.94 Mb), then split and mapped in reverse orientation to the beginning of the chromosome (from 0-5 Mb). By contrast, in the prairie individual, a single contig mapped continuously to the reference genome in this region (37-41.3 Mb) (Figure 3E). Using the long-read sequencing, we localized the inversion breakpoint to basepair resolution (Figure 3F) and found that it lies within a highly repetitive region (Figure S4). Since we found no other forest-specific rearrangements in this region, chromosome 15 likely harbors a simple inversion from 0 to 40.94 Mb. Finally, we used putative centromere-associated sequences in *Peromyscus (16)* to determine that the chromosome 15 centromere is located outside the inversion (Figure 3G). Together, these approaches identified a 40.9-Mb paracentric inversion that is segregating between wild forest and prairie populations and is strongly associated with ecotype differences.

## The inversion is a major region of genetic divergence between ecotypes

To understand the role of the inversion in ecotype divergence, we investigated the frequency and genetic differentiation of the inverted region. First, using genetic principal component analysis (PCA) and levels of heterozygosity, we genotyped the chromosome 15 inversion in wild-caught mice (whole genome re-sequencing: n = 15 forest, 15 prairie) and found that it is nearly fixed for opposite alleles in wild forest and prairie populations (Figure 4A, Figure S5). Second, we compared maximum likelihood-based trees built from the region of chromosome 15 that contains the inversion (*affected* region: 0-40.9 Mb) and the rest of the chromosome (*unaffected* region: 40.9-79 Mb) using RAxML (*17*). In the affected region, forest and prairie mice cluster into two genetically distinct groups based on genotype at the inversion (Figure 4B); this is in contrast to the unaffected region, where mice cluster by ecotype (Figure 4C). Finally, relative genetic differentiation between ecotypes in the affected region, as measured by F_ST_, is more than seven times higher than the genome-wide average (affected region: mean F_ST_ = 0.373; genome-wide average: F_ST_ = 0.052), consistent with the significantly elevated absolute genetic divergence (*d_XY_*) in this region (affected region: mean *d_XY_* = 0.015; unaffected region: mean *d_XY_* = 0.011) (Figure S6). Thus, between ecotypes, the inversion maintains a set of highly divergent sites spanning megabases, including causative loci for both tail length and coat color variation, together in genetic linkage.

**Fig. 4.**
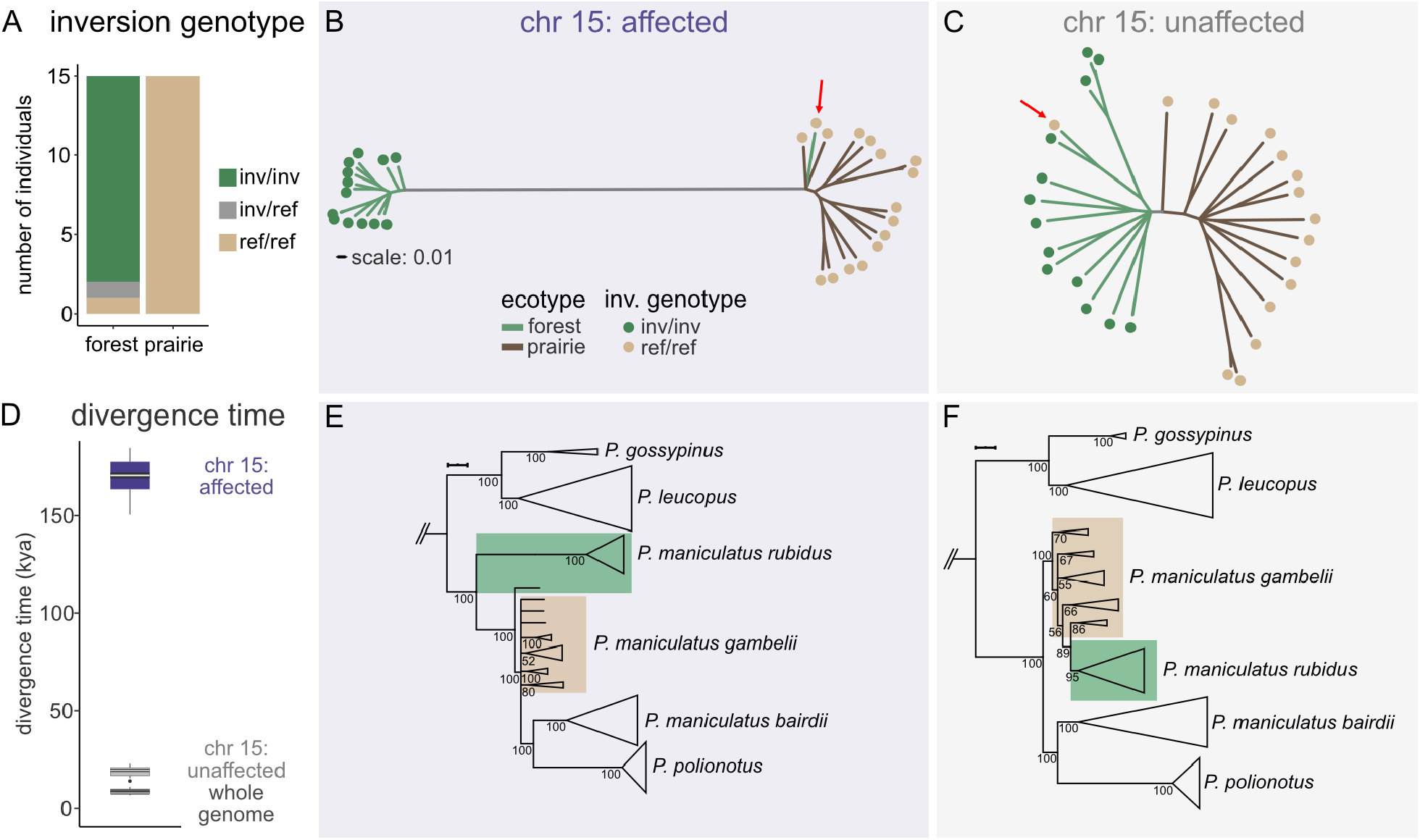
Evolutionary history of the inversion. (A) Frequency of inversion genotype in wild-caught forest (n = 15) and prairie (n = 15) mice (inv/inv, green = homozygous for inversion allele; inv/ref, gray = heterozygous; ref/ref, tan = homozygous for reference allele). (B-C) Maximum likelihood trees for affected (B) (chr15: 0-40.9 Mb) and unaffected (C) (chr15: 40.9-79 Mb) regions of chromosome 15, shown on the same scale. Branch colors indicate ecotype (green = forest; brown = prairie) and dots indicate inversion genotype (tan = homozygous reference, n = 15; green = homozygous inversion, n = 14; heterozygous mouse excluded). Red arrows highlight the forest mouse homozygous for the reference allele. (D) Estimated divergence times between forest (n = 13, only mice homozygous for inversion) and prairie (n = 15) mice for the affected region (purple) and unaffected region (light gray) of chromosome 15, and for all of the autosomes (see supplement for masking strategy) including all mice (dark gray, forest: n = 15; prairie: n = 15). Divergence times (in thousands of years ago, kya) were estimated using SMC++ with 3 generations per year. (E-F) Maximum likelihood trees of *Peromyscus* species for affected (E) (chr15: 0-40.9 Mb) and unaffected (F) (chr15: 40.9-79 Mb) regions of chromosome 15. Trees are rooted with *P. californicus,* not shown. Branches with < 50 bootstrap support are collapsed. Height of triangles is proportional to the number of mice in clade (*P. californicus,* n = 2; *P. gossypinus,* n = 2; *P. leucopus,* n = 22; *P. maniculatus rubidus,* n = 14 in (E), n = 15 in (F); *P. maniculatus gambelii,* n = 15; *P. maniculatus bairdii,* n = 17; *P. polionotus,* n = 17). Green box highlights forest mice and tan box highlights prairie mice. In (E), the single forest mouse outside of the forest clade is homozygous for the reference allele.

The strong genetic differentiation within the affected region suggested that the inversion might pre-date the divergence between the forest and prairie ecotypes. To investigate the evolutionary history of the inversion, we used SMC++ (*18*) to estimate that the inversion and reference alleles likely diverged about 170 kya (assuming 3 generations per year; Figure 4D). By contrast, forest-prairie divergence in the unaffected region and genome-wide are estimated to be 15 and 10 kya respectively, consistent with the divergence occurring near the end of the last glacial maximum approximately ~12 kya. This order-of-magnitude difference in estimated divergence time led us to explore the evolutionary history of the inversion across closely related species, by building phylogenetic trees from the affected and unaffected regions of chromosome 15 using 5 *Peromyscus* species and 3 *P. maniculatus* subspecies (Figure 4E,F, Figure S7). We found that the inverted and reference alleles likely diverged early within *Peromyscus maniculatus,* before the divergence between the forest and prairie ecotypes but after *P. maniculatus* diverged from closely related species. Thus, we infer that, unlike some other known inversions (*19–21*), it is unlikely that the inverted allele introgressed from another extant species; instead, the inversion likely has a long history of polymorphism within the species.

## Inversion frequency in wild populations is consistent with divergent selection

The inversion’s role in genetic and phenotypic divergence between ecotypes suggests that it may be under divergent selection associated with habitat differences. To test the association among genotype, phenotype and habitat in admixed wild populations, we collected deer mice across a sharp habitat transition between the focal forest and prairie sites and estimated habitat type and mean soil hue within 1 km of each capture site (Figure S8, Figure S9; n = 97 mice from 22 sites, supplemented by 12 additional museum specimens from 2 sites). First, we found that much of the transition in both habitat type and soil hue is localized in a narrow region across the Cascade mountain range (Figure 5A,B): for example, while the forest and prairie sites are separated by 500 km, about half of the estimated change in soil hue occurs across just 50 km (10% of the total distance) at the Cascades. Next, phenotypic clines (Figure 5C,D) estimated using either all wild-caught individuals or only those from the central Cascades transect (insets) both identified sharp transitions in tail length and coat color that co-localize with this environmental transition. Specifically, mean tail length changes by 12 mm (45% of the forest-prairie difference) and mean hue changes by 3.2° (60% of the forest-prairie difference) across the same 50-km region. Finally, we found that the inversion frequency decreases from 100% to 62.5% in the central 50 km and then drops to 4% within the next 100 km (i.e., inversion frequency drops from 100% to 4% over less than one third of the total distance between sites; Figure 5E). Together, these data indicate that most of the change in both phenotype and inversion frequency occurs across a sharp environmental transition.

**Fig. 5.**
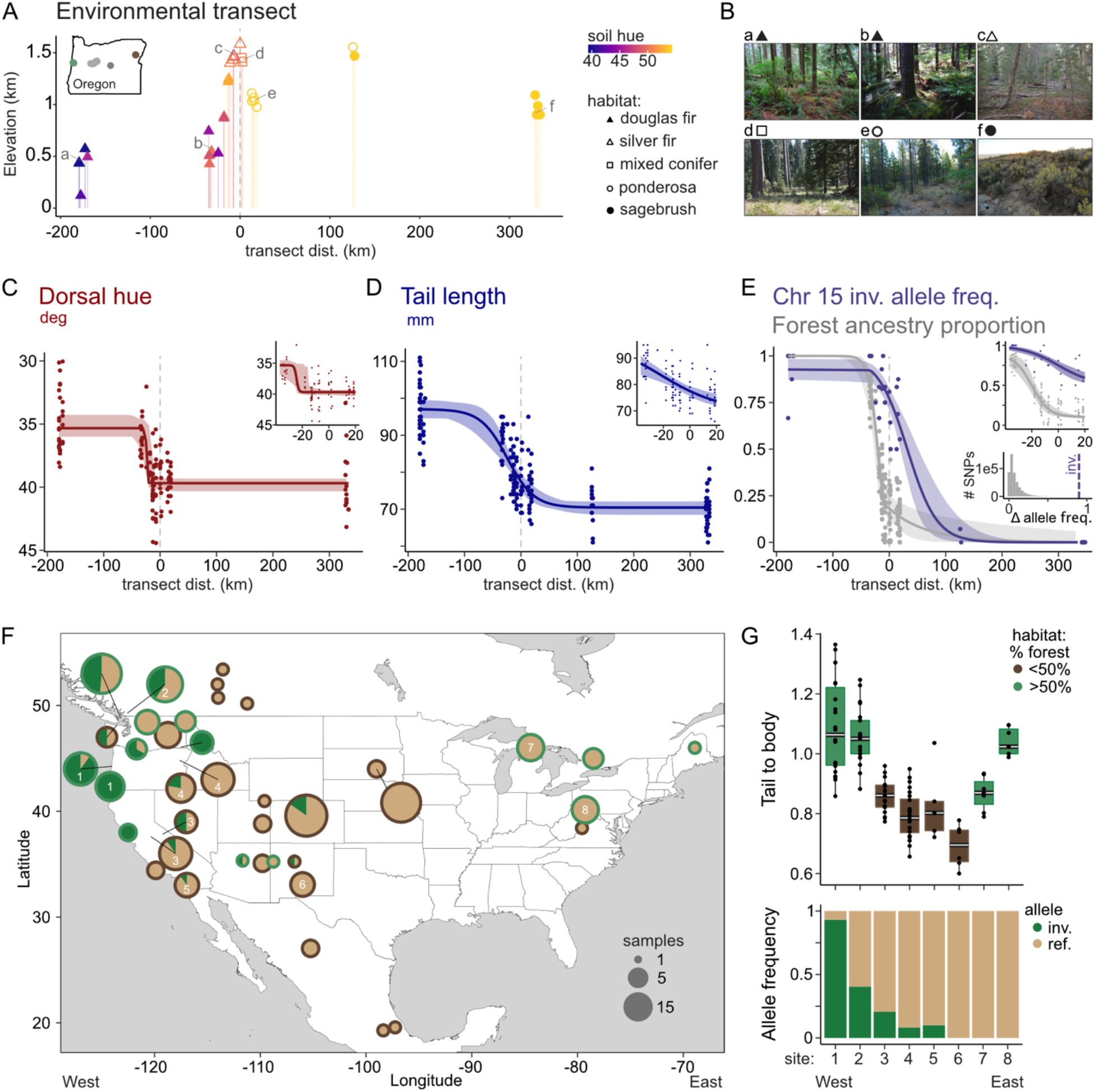
Associations among genotype, phenotype and environment in wild mice. (A) Elevation and habitat characteristics (color = mean soil hue, shape = majority habitat category) at sites across an environmental transect. Letters indicate sites shown in (B). Soil hue and habitat category were estimated for a 1-km radius around each site. Inset: Sampled sites across Oregon, including the forest (green) and prairie (brown) sites, the central Cascades transect (light gray), and additional museum samples (dark gray). Transect distance = east-west distance from the highest-elevation site; dotted lines in A,C,D,E indicate distance = 0. (B) Photos of select capture sites from each habitat type. (C-E) Best fit clines for dorsal hue (C), tail length (D), and genotype (E) fit to the full dataset, with 95% confidence intervals. Insets show best-fit clines using only data from the central Cascades transect. (E) Whole genome ancestry (gray) estimated using ngsAdmix excluding the inversion; points show the ancestry proportion for each individual (gray) and the average inversion frequency for each site (purple). Bottom inset: allele frequency difference (Δ allele freq.) between forest and prairie populations for the inversion (dashed line) and for SNPs used in ancestry estimation (gray). (F) Allele frequency (fill color) for the inversion (green) and reference (tan) alleles at sites across North America for *P. maniculatus* mice (n = 281). Outline color indicates the habitat classification within 1 km of each capture site (> 50% forest: green; < 50% forest: brown). Points are scaled by the number of mice at each site. (G) Tail to body length ratio for *P. maniculatus* populations from across North America (top) with allele frequency of the inversion in green, and reference in tan for each population (bottom). Population numbers correspond to numbered labels on map in (F).

To compare the clinal distribution of the inversion with patterns in the rest of the genome (*22*), we estimated genome-wide forest ancestry proportion using ngsAdmix (best fit: k = 2; (*23*)) and found evidence for admixture between the forest and prairie populations (Figure S10). The estimated forest ancestry proportion changes steeply (from 80% to 14% average forest ancestry) at the environmental transition in the 50-km central region of the transect (Figure 5E). However, the differences in allele frequencies between forest and prairie populations at SNPs outside the inversion are small (Figure 5E; ngsAdmix SNPs have mean allele frequency change of 8%, with 95% of SNPs < 23%). Thus, the steep change in inferred ancestry proportion results from coincident small frequency changes across many loci (Figure S10), in contrast to the steep inversion cline that reflects a large frequency change (90%) between forest and prairie populations at a single locus. The allele frequency difference at the inversion is extreme: it is greater than the maximum difference in allele frequency in 99.92% of similar-linkage-disequilibrium blocks (Figure S11). In addition, we found that the shift in inversion allele frequency is significantly farther east than the change in genome-wide ancestry (assessed using the likelihood profile method (*24*); ML_sum_ = −16.9, ML_comp_ = – 58.2, χ^2^ = 41.4, p = 1.3e-10). We note that the genome-wide cline position is consistent with the mountain range acting as a partial barrier to gene flow. Finally, mixed model analyses using the admixed mice from the central Cascades transect support associations between both inversion genotype and genome-wide ancestry proportion with tail length and hue (Figure S12). Together, the difference in magnitude and location of the inversion frequency shift compared with sites throughout the rest of the genome is consistent with divergent selection favoring different inversion alleles in the forest and prairie habitats.

## Parallel forest phenotypes evolved through partially distinct genetic mechanisms

Previous work reported that the forest ecotype has evolved multiple times in North America (*9*), raising the question of whether the chromosome 15 inversion contributes to adaptation in these independent forest replicates. Using three published datasets (*9, 25, 26*), we determined that the inverted allele is widespread within *P. maniculatus*, particularly in western North America (Figure 5F). To test for an association between the inversion and habitat across the species range, we first characterized habitat in a 1-km radius around each reported capture site. Next, using mixed-effect models implemented in EMMAX (*27*) to control for genetic relatedness among populations, we tested the effect of the inversion as a single locus and found that it is widely associated with both forested habitat (p = 8e-20) and with longer tails (p = 4e-7). Surprisingly, despite this strong association and strikingly similar changes in tail vertebrae (i.e. changes in both vertebrae number and vertebrae length) in eastern and western forest populations (*9*), the inversion was completely absent from eastern forest populations (Figure 5G). These results suggest that the inversion may have played a key role in local adaptation in many, but not all, forest populations; therefore, the genetic architecture that underlies these parallel adaptations is at least partially distinct even within the same species.

## Discussion

Theoretical models suggest that inversions can facilitate local adaptation if they reduce recombination between multiple locally adaptive alleles, even in the absence of epistasis (*28–31*). There is growing empirical evidence supporting the role of inversions in local adaptation (e.g. (*32–38*)), and a limited number of studies have identified inversions associated with multiple distinct traits (e.g. mating types: (*39–41*)). However, few studies have identified inversions that are linked to multiple traits in the context of local adaptation (but see (*42*)). Our results provide new evidence from mammals in support of these models: we found that an inversion maintains variation associated with at least two traits – tail length and coat color, which involve largely distinct developmental and genetic mechanisms – in strong linkage disequilibrium.

Despite gene flow between forest and prairie ecotypes, the inversion ensures that longer tails and darker coat colors are co-inherited, which likely provides a selective advantage in forested habitats.

If inversions are maintained as polymorphisms, they can serve as a source of genetic variation from which a species can adapt to novel or changing environments (*32*), but strong negative fitness consequences of inversions in heterozygotes (e.g. when crossing over produces unbalanced gametes) make this unlikely. In deer mice, however, recombination in terminal inversion polymorphisms is known to be suppressed (*16, 43, 44*), minimizing potential deleterious effects. Indeed, our results suggest that the chromosome 15 inversion was segregating in deer mice long before the forest-prairie ecotypes evolved: the inversion is found at intermediate frequencies in many populations and likely originated within *P. maniculatus* more than 150k years before the last glacial retreat established the modern habitat distributions. Thus, this study highlights how inversion polymorphisms may provide the genetic material for adaptation into newly available habitats. Still, it remains unknown when the tail length and coat color mutations arose relative to the inversion that links them together; future work identifying the causal mutations within the inversion will help elucidate a more precise model of inversion establishment and spread (*45*).

Populations often share standing genetic variation and exchange alleles through gene flow; thus, parallel phenotypic divergence within a species is often due to shared genetic mechanisms (*46–48*). Despite the old origin of the inversion, its large effect on multiple forest traits and its widespread distribution in western forest mice, we find the inversion is absent in eastern forest populations with remarkably similar morphology. Thus, our results suggest that while inversion polymorphisms can be an important source of genetic variation for rapid adaptation, even within a species, distinct genetic mechanisms can result in parallel phenotypes.

In sum, one hundred years after Sturtevant first published his discovery of inversions in laboratory stocks of *Drosophila (49)* and separately, forest-prairie ecotypes were first described in deer mice (*3*), we found that a large chromosomal inversion is key to ecotype divergence in this classic mammalian system, underscoring the important and perhaps widespread role of inversions in local adaptation.

## Acknowledgments

M. Omura, J. Chupasko, M. Mullon, and J. Mewherter helped prepare and accession museum specimens. K. Pritchett-Corning and S. Griggs-Collette provided critical assistance with field work and establishing breeding colonies, and Harvard’s Office of Animal Resources provided assistance with animal care. G.J. Kenagy, D.S. Yang, and E. Kingsley provided invaluable advice at the start of the project, and N. Edelman, A. Kautt, and E. Kingsley provided feedback on the manuscript. We thank C. Lewarch for assistance in the field; S. Niemi, M. Streisfeld, S. Stankowski, G. Binford, S. Bishop, K. Saunders, S. Finch, and T. Schaller for help with field logistics; N. Hughes for assistance with both lab and field logistics; and P. Audano for advice on long-read sequencing and analysis. The University of Washington Burke Museum (UWBM) and the University of New Mexico Museum of Southwestern Biology (MSB) provided specimens used in this study.

## Funding

This work was partially funded by Putnam Expedition Grants to ERH, a Grant-in-Aid of Student Research to ST, and the Chapman Fellowship for the Study of Vertebrate Locomotion to ERH and JTG from the Harvard University Museum of Comparative Zoology, as well as funding from American Society of Mammalogists Grants-in-Aid of Research to ERH and OSH, the Harvard College Research Program to ST, and a Society for the Study of Evolution R.C. Lewontin Early Award to OSH. ERH was supported by an NIH Training Grant to Harvard’s Molecules, Cells, and Organisms graduate program (NIH NIGMS T32GM007598) and by the Theodore H. Ashford Fellowship. OSH was supported by a National Science Foundation (NSF) Graduate Research Fellowship, a Harvard Quantitative Biology Student Fellowship (DMS 1764269), and the Molecular Biophysics Training Grant (NIH NIGMS T32GM008313).

## Author contributions

Conceptualization: ERH, OSH, TBW, ST, and HEH. Formal analysis: ERH, OSH, and TBW. Investigation: ERH, OSH, TBW, ST, JTG, SM, BN, and KMT. Writing: ERH, OSH, TBW, HEH. Visualization: ERH, OSH, TBW. Supervision: HEH. Funding acquisition: HEH.

## Competing interests

Authors declare no competing interests.

## Data and materials availability

Associated data is available as Supplementary Data files S1-S9 and on NCBI Sequence Read Archive (PRJNA687993 and PRJNA688305).

## Notes

https://www.ncbi.nlm.nih.gov/bioproject/PRJNA687993

https://www.ncbi.nlm.nih.gov/bioproject/PRJNA688305

### Competing Interest Statement

The authors have declared no competing interest.

