## Supplemental Materials for "A chromosomal inversion drives evolution of multiple adaptive traits in deer mice"

**This PDF file includes:**

Materials and Methods
Figs. S1 to S18
Tables S1 to S4
Captions for Data S1 to S9

**Other Supplementary Materials for this manuscript include the following:**

Data S1 to S9:
- Data S1: Wild-caught specimens from this study
- Data S2: *Peromyscus* phylogeny sample ids and museum collections
- Data S3: Genotypes and phenotypes for mice from across North America
- Data S4: Wild-caught forest, prairie and transect mice phenotypes
- Data S5: Lab-born forest and prairie body dimensions
- Data S6: Lab-born forest and prairie coat color phenotypes
- Data S7: F2 hybrid phenotypes
- Data S8: Habitat category by site across transect
- Data S9: Soil color by site across transect

### MATERIALS AND METHODS

#### Sampling and measuring forest and prairie mice

Field sampling

To test for phenotypic differences between forest and prairie ecotypes, we captured deer mice (*Peromyscus maniculatus*) from two populations separated by approximately 510 km: 32 adult mice in sagebrush steppe in eastern Oregon ('prairie' ecotype) and 44 mice (39 adult) in temperate rainforest in western Oregon ('forest' ecotype); of these, we used 20 mice per population to establish laboratory colonies (see below). In addition, we sampled 136 mice (98 adult) from 22 sites focused in a 50 km transect across the Cascade mountain range, between the forest and prairie populations, with 1-19 mice captured at each site. We included both subadult and adult mice for genetic analysis but analyzed phenotypes from adults only. Trapping took place in September-October 2015 and August-September 2016 using Sherman live traps. With a few exceptions, the wild-caught mice were ultimately accessioned in Harvard's Museum of Comparative Zoology Mammal Collection (see Data S1).

Establishing laboratory colonies

To establish laboratory colonies, we transported and quarantined 40 wild-caught mice (20 male, 20 female) at Harvard University. From these founders, we maintained two separate laboratory colonies, representing the forest ecotype, *P. m. rubidus* (4 original productive breeding pairs) and the prairie ecotype, *P. m. gambelii* (7 original productive breeding pairs). After quarantine, we housed mice at 23°C on a 16h:8h light:dark cycle in standard mouse cages (Allentown Inc, Allentown, NJ, USA) with corncob bedding (The Andersons, Inc, Maumee, OH, USA), cotton nestlet (Ancare, Bellmore, NY, USA), Enviro-Dri (Shepherd Specialty Papers, Watertown, TN, USA), and either a red tube or a red hut (BioServ, Flemington, NJ, USA). We provided animals with *ad libitum* water and mouse chow (LabDiet Prolab Isopro RMH 3000 5P75). We used the HAN rotation breeding scheme (*1*, *2*) to maintain outbred colonies.

Morphological measurements

For each individual, we took standard morphological measurements while the animal was alive (for colony-founding individuals) or immediately following euthanasia. We measured total length (nose to tail tip), hindfoot length, ear length, and tail length, and weight. We calculated body length as the difference between total and tail length. Laboratory-born mice were measured as adults, between 60 and 70 days old.

Pigmentation measurements

To measure coat color, we used a FLAME UV-VIS spectrometer with a pulsed xenon light source, a 400 um reflectance probe, and OceanView software (Ocean Optics) to measure 3-5 reflectance spectra from each of 3 body regions (dorsal stripe, flank, and ventrum). We used a custom R script (available at github.com/emilyrhager/HagerHarringmeyer) to obtain brightness, hue, and saturation values in the range 400-700 nm with 1 nm bin width, using the segment classification approach (*3*) with formulae as described for CLR v 1.05 (*4*). For every trait, we calculated the median value for each body region and individual. We measured forest and prairie mice shortly after euthanasia (for wild-caught mice) or while frozen (for laboratory-born specimen). For the transect analysis, all wild-caught mice were fixed in formalin, stored in ethanol, and then air-dried before measuring the dorsal and flank regions with the spectrophotometer.

Statistical analysis of phenotypes

To test for differences between the forest and prairie subspecies, we used t-tests for all 14 morphological and pigment traits (body length, weight, tail length, ear length, and hindfoot length; and brightness, hue and saturation in the dorsal, flank and ventral regions). We corrected all results for multiple testing using the Bonferroni-Holm method, and performed statistical analysis using R v 3.6 (*5*).

**Forward genetic mapping**

F2 intercross design

We conducted a reciprocal cross between laboratory-raised 'forest' *P. m. rubidus* and 'prairie' *P. m. gambelii* to generate F1 hybrids, and then intercrossed sibling F1 hybrids to generate F2 animals. Based on the measurements of laboratory-reared mice, we used R/qtl (*6*) to estimate that n = 500 hybrids would provide 80% power to detect loci explaining ~5% of the variance in tail length; we therefore tested 555 F2 hybrids in total (forest female x prairie male, n = 203 F2s from 12 F1 breeding pairs; prairie female x forest male, n = 352 F2s from 13 F1 breeding pairs). All 555 F2 hybrids descended from four individuals (the cross founders: one sibling male and female of each subspecies) and were measured for morphological traits as described above before genotyping.

Ancestry assignment in F2 hybrids

To genotype F2 hybrids, we generated low-coverage Double Digest Restriction Associated DNA (ddRAD) sequencing libraries (*7*) and then assigned hybrid genotypes with the Multiplexed Shotgun Genotyping (MSG) pipeline (*8*), using high-confidence fixed variants identified from whole genome sequencing of the cross founders.

*ddRAD sequencing and joint genotyping of F2 hybrids*

Briefly, we extracted DNA from liver tissue using the Autogenprep 965 (Autogen), digested the DNA with restriction enzymes MluCI and NlaIII (New England Biolabs), and then ligated adapter sequences (MluCI overhang: individually barcoded; NlaIII overhang: common, biotinylated). We pooled samples in groups of no more than 48 samples and used a Pippin Prep (Sage Science) to select fragments between 216 and 276 bp. We enriched for DNA with both adapters using streptavidin beads (Dynabeads, Invitrogen) and then performed 9 cycles of PCR amplification with Phusion Taq polymerase, adding pool-specific adapter sequences. After performing quality control with TapeStation (Agilent) and measuring concentration with Qubit 3.0 (ThermoFisher) and qPCR, we combined all pools into a single library and sequenced 125 bp paired-end reads across four lanes of Illumina HiSeq v4. To avoid clustering problems during sequencing that can be caused by shared overhang sequence across ddRAD reads, we included a previously-generated RNAseq library as a diversity spike-in (10% of each lane). We performed ddRAD sequencing for all 555 F2 hybrids as well as the 4 cross founders and 49 F1s. Sequencing was performed at The Bauer Core Facility at Harvard University and computations were performed using the Harvard Research Computing cluster.

We demultiplexed reads based on individual barcodes, and then mapped sequencing reads to the *P. maniculatus bairdii* reference genome (NCBI accession: GCA_003704035.3) using BWA-MEM, with –p to indicate interleaved paired-end fastq input, and –M to mark short split hits as secondary for compatibility with Picard. The median read depth was 635,012 mapped reads per individual and we excluded the F2 hybrids (n = 8) with read depth less than 75,000 total mapped reads. From the mapped bam files, we ran *HaplotypeCaller* (GATK3.8) with the default heterozygosity prior (-hets = 0.001) and –ERC GVCF to produce per sample gVCFs. Then, we ran *GenotypeGVCFs* (GATK3.8) to jointly genotype the samples. We performed hard filtering of SNPs based on GATK best practices (filtering variants with QD < 2.0, FS > 60.0, MQ < 40.0, MQRankSum < -12.5, ReadPosRankSum < -8.0) using *VariantFiltration* (GATK3.8).

*Identifying fixed SNPs between the forest and prairie cross founders*

To identify fixed variants between the forest and prairie cross founders, we performed whole-genome re-sequencing of the 4 laboratory-born cross founders (2 forest *P. m. rubidus* and 2 prairie *P. m. gambelii*), as well as 3 of the 4 wild-caught parents of these founders (both parents of the forest founders and the female parent of the prairie founders). We extracted DNA from ~20mg of liver tissue and generated sequencing libraries using a PCR-free KAPA HTP kit. Following enzymatic fragmentation, we used size selection to enrich for a 450 bp insert size and ligated Illumina adapters. We sequenced the resulting libraries using 150 bp paired-end sequencing on an Illumina NovaSeq S4 flowcell.

Following demultiplexing, we mapped sequencing reads as described above and marked optical and sequencing duplicates using *MarkDuplicates* (Picard), with OPTICAL_DUPLICATE_PIXEL_DISTANCE=2500 to account for artifacts generated from the patterned flowcell found in the NovaSeq S4. To call variant sites, we first used *HaplotypeCaller* (GATK4.1) on each sample as above. Then, intermediate haplotype files for all individuals were consolidated into a GenomicsDB structure using *GenomicsDBImport* (GATK4.1), which was used to create variant + invariant cohort-level vcfs for each chromosome with *GenotypeGVCFs* (GATK4.1). We performed hard filtering of SNPs with parameters as described above, and hard filtering of INDELs with QD < 2.0, FS > 200.0, ReadPosRankSum < -20.0, SOR > 3.0. We also filtered invariant sites with QUAL ≥ 20 using bcftools. Finally, we masked all variant calls with read depth < 5.

To create a set of fixed variants between the prairie and forest founders, we combined the ddRAD and whole genome re-sequencing data using *CombineVariants* (GATK3.8). We then performed SNP filtering independently for each set of reciprocal cross founders. We used *SelectVariants* (GATK3.8) to select SNPs that showed fixed differences between the forest and prairie founders (number of fixed SNPs = 1,022,051 for prairie female x forest male; 984,603 for forest female x prairie male). Since we expected the F1s to be heterozygous at each fixed SNP, we excluded any SNP for which the ratio of the number of reference to alternate calls across all F1 reads fell within the top or bottom $10^{th}$ percentile of a binomial distribution (p = 0.5, n=total number of F1 reads at a given SNP). Finally, we removed SNPs for which a cross founder was called homozygous for one allele, but the founder's parent was homozygous for the opposite allele. Post-filtration, we obtained a total of 793,928 fixed SNPs for prairie female x forest male and 784,259 fixed SNPs for forest female x prairie male.

*Ancestry assignment of F2 hybrids with the multiplexed shotgun genotyping pipeline*

We used the hidden Markov model implemented in the Multiplexed Shotgun Genotyping (MSG) pipeline (*8*) to assign ancestry in F2 hybrids following (*9*). In brief, we used samtools *mpileup* to extract the appropriate set of fixed SNPs from the mapped bam files of F2 hybrids, requiring that SNPs were at least 500 bp apart to ensure they were from independent reads. Then, we ran the fit-HMM step of MSG (fit-hmm.R) on the filtered mpileups with the settings: deltapar1=0.1, deltapar2=0.1, rfac=1; priors=0.25,0.5,0.25; theta=1; one_site_per_contig=1; recRate=25. We combined the fit-HMM genotype probabilities across all F2 hybrids using combine.py (https://github.com/JaneliaSciComp/msg/), which interpolates missing genotypes, resulting in a total of 978,808 markers. Next, we thinned the marker density, retaining every $100^{th}$ marker and keeping neighboring markers for which at least one F2 hybrid had genotype conditional probabilities differing by 0.1 using pull_thin (https://github.com/dstern/pull_thin) with difffac=0.1. After thinning, we had a total of 109,356 markers. Finally, we created an R/qtl object using read.cross.msg.1.5.R (https://github.com/dstern/read_cross_msg/), where we directly imported genotype probabilities from fit-HMM as the R/qtl genotype probabilities and added the phenotype data. We dropped markers with missing genotypes for more than 25 individuals, resulting in 108,575 markers used for mapping.

<u>Quantitative Trait Locus mapping</u>

To identify genomic regions associated with morphological and pigment variation after testing for correlations among traits (Figure S13), we performed QTL mapping using the extended Haley-Knott method in R/qtl v. 1.45.11; ('ehk' method in the *scanone* function; (*6*, *10*)). We used permutation tests to obtain thresholds for genome-wide significance, with 1000 permutations for the autosomes and a separate significance threshold for the X chromosome (18,600 permutations), and estimated confidence intervals using the 95% Bayes credible interval (*bayesint* function in R/qtl). To estimate the additive and dominance effect sizes and the percent of the total phenotypic variance associated with each locus, we fit a linear model including the peak markers for all significant loci and any covariates using the *fitqtl* function in R/qtl (method = 'hk', including additive effects of each locus). Because tail and hindfoot length were significantly correlated with body length (tail-body: r = 0.26, 95% CI = 0.18-0.33; hindfoot-body: r = 0.18, 95% CI = 0.09-0.26), we included body length as an additive covariate for these traits. For all traits, we also tested whether there was a statistically significant association with 3 additional possible covariates: age, cross direction, and sex. Where such associations existed (with age, cross direction, and sex for foot length, age and sex for tail length, and cross direction only for pigmentation traits), we ran additional QTL models including these as additive covariates. As including these extra covariates did not substantially alter the results (Table S2), we report the results from the simpler models here.

Recombination breakpoint analysis

We determined locations of recombination breakpoints in the F2 hybrids from their genotype probabilities in the R/qtl object. We localized breakpoints to the midpoint of where an individual's homozygous genotype probability transitioned from below 0.05 to above 0.95 or from above 0.95 to below 0.05. For consecutive breakpoints involving transitions between the same two genotypes, if the breakpoints localized within 500 kb of each other, we excluded the breakpoints from our analysis as such closely neighboring breakpoints were likely due to short tracts of spurious ancestry calls.

**Identifying an inversion on chromosome 15**

In our QTL analyses, we identified a region of chromosome 15 that showed limited recombination over a large physical distance (~40 Mb). To investigate this region further, and in particular, to determine whether the lack of recombination was due to a chromosomal rearrangement, we first used whole genome re-sequencing data in a larger sample of wild-caught mice. Using these data, we tested whether this region was more differentiated between the forest and prairie populations compared to the rest of the genome, consistent with a structural variant inhibiting gene flow in the wild. We then used long-read sequencing to determine the nature of the rearrangement.

Estimating $F_{ST}$ between forest and prairie populations

To assess genome-wide levels of nucleotide variation, we performed whole genome re-sequencing for 30 wild-caught mice (forest n = 15; prairie n = 15). We generated sequencing libraries, called variants, and performed filtering together with the cross founders (see above: "Identifying fixed SNPs between the forest and prairie cross founders"), except that for 4 mice (2 forest, 2 prairie) we generated sequencing libraries using the PCR-based KAPA HTP kit.

We estimated $F_{ST}$ using the program *angsd*, which can take genotype uncertainty into account instead of relying on called genotypes (*11*). First, input bam files were used to generate site allele frequency likelihood files (SAFs) with the following command: "angsd -gl 2 -doSaf 1 -minMapQ 30 -minQ 20 -C 50 -baq 1". We next estimated individual and pairwise site frequency spectra (SFS) using *realSFS fst index* and setting the parameter "-nSites" to 5e8. Finally, we calculated $F_{ST}$ using *realSFS* with the individual and pairwise SFS as priors. Global $F_{ST}$ and sliding window $F_{ST}$ were estimated using "*realSFS fst stats*" and "*realSFS fst stats2 -win 10000 -step 1000*" respectively.

Linkage disequilibrium in forest and prairie populations

We next investigated linkage disequilibrium across chromosome 15 in the forest and prairie mice (n = 30). Using the wild-caught re-sequencing data, we filtered chromosome 15 SNPs to include only biallelic SNPs with < 5% of samples missing genotypes and minor allele frequency > 0.1. We then thinned SNPs to $\leq$ 1 SNP per 100 kb, resulting in a total of 786 SNPs. We used vcftools --geno-r2 to compute $r^2$ between each pair of SNPs, using genotypes to calculate correlations to accommodate unphased data.

Long-read sequencing

To better characterize the putative chromosome 15 rearrangement, we performed long-read sequencing on 2 individuals: one forest mouse homozygous for the structural variant and one prairie mouse homozygous for the reference allele. First, we extracted high-molecular weight (HMW) DNA from 200 uL fresh blood using the MagAttract HMW DNA mini kit (Qiagen), following the Whole Blood protocol (Qiagen), using wide-bore pipette tips to prevent shearing at the elution step. We quantified the resulting DNA using a Genomic DNA ScreenTape on the Tapestation 4200 (Agilent). Library preparations and sequencing were performed at the University of Washington's PacBio Sequencing Core. In brief, SMRTbell libraries were prepared with the SMRTbell Express Template Prep Kit 2.0 (PacBio). We performed a size selection of 30 kb for the forest sample using the BluePippin (Sage Science), and we did not perform any size selection for the prairie sample since total library mass was below 500 ng. Then, we sequenced each on a Sequel II SMRTcell 8M (PacBio), the forest sample with a 15-hour movie and the prairie sample with a 30-hour movie. The unique molecular yield was 131.3 Gb for the forest sample and 134.7 Gb for the prairie sample, with the longest subread N50 of 37,943 bp and 36,619 bp, respectively.

For each sample, we generated *de-novo* assemblies using the program *canu* (*12*). Given the high levels of heterozygosity, we specified the parameters corOutCoverage=200 and correctedErrorRate=0.15 to allow some read mismatch and therefore combine haplotypes, achieving a haploid (rather than diploid) assembly. Contig N50s for the de-novo forest and prairie assemblies were 1.37 Mb and 1.22 Mb, respectively. We then used the program *mummer* to align each de-novo assembly to the *P. maniculatus bairdii* reference genome (*13*). We implemented the *nucmer* command with default parameters to accommodate potential rearrangements between the draft assembly and reference genome.

After determining that chromosome 15 harbors a large chromosomal inversion, we used the program *sniffles* (*14*) to search for additional structural variants on the chromosome in an unbiased manner by calling variants from long-read mapping. From each movie, we converted the subreads bam to fastq using bam2fastx (PacBio). We then aligned fastq files to the *P. maniculatus bairdii* reference genome (Pman2.1.3) using the program ngmlr with the pacbio preset parameter "-x pacbio". Next, we converted output sam files to bam format with *samtools view* and added readgroups with *AddOrReplaceReadGroups* (Picard) for downstream compatibility. For each individual, we called variants using the program *sniffles* with the parameter "-d 5000". We then merged these raw variant calls with the program *SURVIVOR* using the parameters "1000 1 1 -1 -1 -1". This merged callset was then used to re-genotype each individual with *sniffles*, so as to obtain a genotype for each individual at every site. We merged the final callset again using *SURVIVOR* and considered any large structural variants (>100 kb) that were fixed differences between the sequenced forest and prairie mice as a candidate set, which we verified using the contig alignment.

**Determining the frequency of the inverted haplotype**

To genotype the 30 wild-caught individuals for the inversion, we first examined patterns of relatedness and heterozygosity across chromosome 15. Specifically, we calculated heterozygosity for each individual, within the affected (inversion, 0-41 Mb) and unaffected (no inversion, 50-79 Mb) chromosome 15 regions using "*vcftools –het*". Next, we used *plink –pca* to perform PCA on all biallelic SNPs from both regions. Three distinct clusters in PC1 suggested 3 genotypes: homozygous for the inversion, heterozygous, and homozygous for the reference allele. These genotypes were consistent with observations of decreased heterozygosity in homozygous inversion mice and increased heterozygosity in heterozygous mice.

**Phylogenetic analyses**

Chromosome 15 trees: forest and prairie mice

To assess forest-prairie divergence at the inversion, we constructed maximum likelihood trees for the affected (0-41 Mb) and unaffected (41-79 Mb) regions of chromosome 15 for the wild-caught forest (n = 14) and prairie (n = 15) mice, excluding a single forest mouse heterozygous for the inversion. From the whole-genome re-sequencing vcf, we thinned SNPs to a maximum of 1 SNP per 100-bp, converted the vcf to a PHYLIP matrix using vcf2phylip.py (*15*) (https://github.com/edgardomortiz/vcf2phylip), and removed invariant sites using ascbias.py (https://github.com/btmartin721/raxml_ascbias). This resulted in 76,163 and 50,909 SNPs for the affected and unaffected regions, respectively. We built trees with RAxML v8.2.12 (*16*) using the GTRCAT model, with the conditional likelihood method, -asc-corr=lewis, to correct for the ascertainment bias due to using SNPs (*17*). We ran 100 bootstraps, with "-f a" to perform rapid bootstrap analysis. We visualized trees in iTOL (*18*) and collapsed branches with bootstrap support < 75%.

Chromosome 15 trees: *Peromyscus* species

To estimate when the inversion arose within the *Peromyscus* genus, we next built maximum likelihood trees for the chromosome 15 affected and unaffected regions across multiple *Peromyscus* species: *P. californicus* (n = 2), *P. gossypinus* (n = 2), *P. leucopus* (n = 22), *P. polionotus* (n = 17), *P. maniculatus bairdii* (n = 17), *P. maniculatus gambelii* [prairie] (n = 15), *P. maniculatus rubidus* [forest] (n = 15); sample sources are given in Data S2. Whole genome re-sequencing and joint genotyping was performed for all samples as described above. From the whole-genome re-sequencing vcf, we thinned SNPs to a maximum of 1 SNP per 100-bp, and prepared the SNP set as described for forest-prairie trees. This resulted in 117,848 and 98,113 SNPs for the affected and unaffected regions, respectively. We built trees with RAxML (v8.2.12, (*16*), using the GTRCAT model, with the conditional likelihood method and bootstrapping as described above. We rooted trees with *P. californicus* and visualized trees in iTOL, collapsing branches with bootstrap support < 50%.

**Nucleotide diversity across chromosome 15**

To investigate genetic diversity across the affected and unaffected chromosome 15 regions, we used the PopGenome package in R (*19*). We imported biallelic SNPs from the whole-genome re-sequencing vcf of the wild-caught forest (n = 13) and prairie (n =

15) mice into PopGenome, excluding the 2 forest mice carrying at least one reference allele. We computed statistics in 10-kb windows, with step size of 10 kb. We also calculated $d_{XY}$ using the *diversity.stats.between* function and pairwise nucleotide diversity separately for the forest and prairie mice using the *diversity.stats* function.

**Inversion age and population divergence time estimates**

To estimate the age of the inversion relative to the forest-prairie ancestor, we used the Sequential Markov Coalescent + Plenty of Unlabeled Samples (SMC++) (*20*) method on the re-sequencing data from the 30 wild-caught mice. This approach combines information from the distribution of coalescence times in one or more whole genomes and the site-frequency spectrum (SFS) in the rest of the sampled population. For our implementation, we estimated ages for 3 datasets: (1) affected region: chr15: 0-41 Mb, forest mice homozygous for the inversion (n = 13) and all prairie mice (n = 15); (2) unaffected region: chr15: 50-79 Mb, forest mice homozygous for inversion (n = 13) and all prairie mice (n = 15); (3) all autosomes: all forest and prairie mice (n = 30). For each dataset, we followed three steps. First, we used *vcf2smc* to convert a vcf of biallelic SNPs to the SMC++ input format. In this conversion step, we specified the highest-coverage sample as the "distinguished individual" (option -d) and provided a mask for low-mappability regions following the SNPable protocol (http://lh3lh3.users.sourceforge.net/snpable.shtml) and for $\geq$50% missing genotypes at a given site (variant or invariant). Second, we used the *estimate* command to estimate per-population demographic histories, with the input mutation rate of 5.3e-9 (*21*) and the following parameters: '--timepoints 1e3 5e7 --Nmax 1e8 --spline cubic'. Third, we ran the identical command on 20 bootstrapped SMC++ input files created with a custom script. The models from these individual population histories (Figure S14) as well as pairwise SMC++ input files for the combined forest and prairie populations (normal and bootstrapped) were used as input for *split* to estimate the time since population divergence. *Split* was run with default parameters. Finally, we converted time estimates to years, assuming 3 generations per year (*22*).

**Distribution of the chromosome 15 inversion and morphological traits across a transect**

<u>Cascades transect sampling</u>

To test for a relationship among phenotypic variation, environmental variation, and allele frequency change in the wild, we sampled additional mice (n = 136) between the forest and prairie populations. Specifically, we focused sampling in a 50-km region running east-west across the Cascades mountain range for several reasons: first, the Cascades represent a sharp habitat transition from wetter, coastal forest to dry interior forest, and also form a contact zone for many other species and subspecies ((*23*–*26*) among others), and second, our initial (2015) samples indicated a sharp phenotypic change at this location, which recapitulated most of the difference between the forest and prairie mice. In addition to sampling across the Cascades, we also included museum specimens (n = 12) from a site intermediate between the Cascades samples and the eastern-most prairie site. For these museum specimens, we were able to obtain comparable data for tail length and inversion genotype (see below) but not for pigmentation or whole genome ancestry.

Habitat characterization

To determine whether phenotypic and/or allele frequency change was associated with environmental variation, we used QGIS v. 3.4 (*27*) to evaluate vegetation type and soil color across the full transect. To assess soil color, we used the publicly available STATSGO2 data from USGS (*28*). To estimate local soil characteristics at each site, we found the total area within a defined radius of the trapping location (i.e. 0.5, 1, or 2 km) that belonged to each official USDA soil series and generated a weighted average of the Munsell soil color, value and chroma. We used a custom python script (with results verified by manual review) to gather Munsell color characterizations for the top-most layer of each soil series from the USDA Official Soil Series Descriptions (*29*). If the top-most layer did not have Munsell color information, we used the values from the top layer that did. We used the Munsell characteristics for moist soil if the description listed the series as ‘usually moist’, udic or aquic moisture regimes, or dry less than 90 consecutive days in summer; otherwise, we used the dry soil characteristics (i.e., if the series was described as ‘usually dry’, aridic moisture regime, or dry more than 80 days in the summer). For analysis, we converted Munsell hues to degrees with hue 5R at 0°; thus, 5YR = 36°, 7.5YR = 45°, 10YR = 54°, and 2.5Y = 63°. Finer-scale soil survey (i.e. SSURGO) data were not available for the central Cascades transect, but where such surveys were available, the estimated soil characteristics we obtained from the two datasets matched. Because the results were similar across all three radii (Figure S9), we chose to focus on the 1-km data. For vegetation data, we made use of detailed habitat models from the Oregon Biodiversity Information Center (Oregon Spatial Data Library, https://spatialdata.oregonexplorer.info/) (*30*) to calculate the proportion of area within the same three radii around the sampled sites occupied by each habitat type. These results also did not depend heavily on the chosen radius (Figure S8). To display maps, we used ggmap (*31*).

Genotyping the inversion and estimating ancestry in Cascades mice

*Whole genome re-sequencing and joint genotyping*

We performed low-coverage whole-genome re-sequencing of the mice (n = 136) from the 22 sites in the central Cascades transect. We extracted genomic DNA from liver tissue using proteinase K digestion followed by the Maxwell RSC (Promega) DNA extraction. For library preparation, we used the Nextera XT kit with Illumina adapters, performing the reactions at 1/4 volume. We quantified libraries with the TapeStation (Agilent), pooled the samples into a single library, and quantified the pooled library by qPCR. We then sequenced the library on a full flow cell of Illumina NovaSeq SP with paired-end sequencing of 150 bp reads.

After de-multiplexing reads based on the Illumina barcodes, we mapped the reads from the fastq files to the *P. maniculatus bairdii* reference genome following the protocol described above. The median read depth for these samples was 15,347,998 mapped reads per sample, which corresponds to ~1.5X sequencing coverage across the genome. From the mapped bam files, we created cohort level vcfs as described above.

*Determining inversion genotypes*

To determine chromosome 15 genotypes, we created a set of fixed SNPs to differentiate the inversion and reference haplotypes. We used *SelectVariants* (GATK3.8) to select fixed SNPs in the affected region between the wild-caught mice homozygous for the inversion (n = 13 forest) and those homozygous for the reference allele (n = 15 prairie, n = 1 forest). We filtered this set of SNPs by requiring that the one forest mouse heterozygous for the inversion must be heterozygous at the selected SNPs, resulting in a set of 37,242 SNPs fixed between the inversion and reference haplotypes. We then performed the fit-HMM step of MSG for the Cascades mice using this set of SNPs for the affected region, as above.

For the additional museum specimens (n = 12), we extracted genomic DNA from liver tissue using proteinase K digestion followed by the Maxwell RSC. Then, we used four custom Taqman SNP genotyping assays (Life Technologies) to genotype diagnostic SNPs between the inverted and reference haplotypes (Table S3). All genotyping reactions were performed with 1-10 ng of genomic DNA, using the following cycling parameters: 95 °C for 10 minutes followed by 40 cycles of 95 °C for 15 s, 60 °C for 1 minute. For all individuals, the four assays gave the same genotype results.

*Ancestry estimates*

As the forest and prairie populations had low genetic differentiation across the genome, we used ngsAdmix (*32*) to estimate ancestry proportions for each Cascades transect individual (n = 136), and for the previously sequenced wild-caught forest and prairie individuals (n = 30). To identify a set of SNPs for ngsAdmix, we selected biallelic SNPs from all autosomes that had at least 50% of all samples (forest, prairie, and Cascades mice) with non-missing genotypes using *SelectVariants* (GATK3.8) and thinned SNPs to be at least 1 kb apart to avoid SNPs in strong linkage disequilibrium (Figure S15). We excluded the affected region so that whole-genome ancestry estimates were not influenced by the inversion genotypes, resulting in a total of 472,692 SNPs. To run ngsAdmix, we created a beagle file using a custom R script to convert GATK PLs to genotype likelihoods. Then, we ran ngsAdmix with minMAF=0.1 and k=1-6. We performed 20 ngsAdmix runs per k, each with a different random seed. Using CLUMPAK (*33*, *34*), we determined that k=2 was the best number of clusters (Figure S10A). Thus, we ran ngsAdmix with minMAF=0.1 and k=2 for estimating ancestry coefficients. We determined confidence bounds for the ancestry estimates by performing 100 iterations of subsampling the input SNP set to 25% and re-running ngsAdmix (Figure S10B).

In addition, we used ngsAdmix to confirm MSG genotypes for the inversion. From the set of thinned SNPs as described above, we had 7,090 SNPs in the affected region of chromosome 15. Using this set of SNPs, we ran ngsAdmix as described above, and the ngsAdmix ancestry estimates confirmed MSG genotypes for the inversion (Figure S10F). Finally, we determined the allele frequency difference between the forest and prairie focal populations for all SNPs used in ngsAdmix.

*Fitting clines to genotypes and phenotypes*

First, to test whether mouse pigmentation was associated with soil color and value, we calculated the correlation between mouse hue and mean soil hue at the site of capture, between mouse brightness and mean soil value, and between mouse saturation and mean soil chroma (Figure S16).

To determine spatial changes in morphological and genetic variation, we next used R to fit sigmoid clines for each trait separately using both the full dataset (i.e., wild-caught forest, prairie, and Cascades mice, with additional museum specimens for tail length and the inversion) and, separately, data restricted to the Cascades. We first converted the spatial coordinates of each site into an east-west transect distance, with the central point (distance = 0 km) at the highest elevation. We then used the *distRhumb* function from the package geosphere (v. 1.5.7, (*35*)) to calculate east-west distances between this central point and each other sampled site.

We next used the package HZAR v. 0.2.5 (*36*) to fit clines for tail length, dorsal and flank hue, chromosome 15 inversion genotype, and whole-genome ancestry. For phenotype values, we fit 5 cline models, which varied only according to whether and how exponential tails were fit (tails = 'none', 'left', 'right', 'mirror', and 'both'); exponential tails indicate stepped clines (*37*). For genotype data, we fit 10 cline models, which varied by how exponential tails were fit (tails = 'none', 'left', 'right', 'mirror', and 'both') and by the scaling of minimum and maximum allele frequencies (scaling = 'fixed' for minimum and maximum fixed as minimum and maximum of observed mean data, 'free' for minimum and maximum as free parameters). We selected the best model for each trait and genotype using AICc values. We bounded the values for center and width such that the center could not be more than 10 km outside the sampled region, and the width no more than the total transect distance + 20 km (*36*). For phenotypic data, we also bounded the variance in the center of the cline ('varH') to be no more than 1.5x the total variance in the dataset, which helped runs to converge. For each model, we ran 3 independent chains (varying the random seed) with 3 runs each, with chain length 1e6, thin 1e3, and burnin 1e5 for phenotype data, and chain length 1e5, thin 100, and burnin 1e4 for genotype data. We assessed chain mixing by visually inspecting the trace and using the potential scale reduction factor (*gelman.diag* in the coda package, (*38–40*)).

Finally, to test for coincidence of the genome-wide ancestry and chromosome 15 inversion clines, we used the likelihood profile method (*41*). For each variable to compare (i.e. whole genome ancestry proportion and inversion genotype), we constructed likelihood profiles by finding the likelihood of the best-fit cline model with the center fixed at each of 21 values between -30 and 30 km, the region where all best-fit cline centers were located. We fit exponential tails according to the best-fit model from the initial analysis (i.e. tails = 'right' for ancestry, and 'left' for chromosome 15; scale = 'fixed') and used the full, not transect-restricted, dataset. We used a chi-square test with test statistic equal to the sum of the maximum likelihood values for each trait, MLsum, minus the maximum value of the summed likelihood profiles, MLcomp, to test for cline coincidence of particular clines, with degrees of freedom equal to the number of compared traits minus one. If all the clines are co-located, MLsum and MLcomp should be approximately equal, while if not, MLcomp will be significantly less than MLsum.

<u>Linking genotype and phenotype in Cascades mice</u>

To test whether genotype at the chromosome 15 inversion was associated with phenotype in the Cascades mice, we used linear mixed effects models. Specifically, we used tail length, dorsal hue, and flank hue as response variables, with fixed effects of genotype at the chromosome 15 inversion and the estimated proportion of genome-wide forest ancestry excluding chromosome 15 from ngsAdmix, and a random effect of site of capture. We compared these 'full' models to models that included ancestry proportion and capture site, and to models that included site alone, using AIC values to select the best model. In the Cascades, sample sizes for the 3 chromosome 15 genotypes were highly variable: in particular, while there were 59 forest homozygotes and 34 heterozygotes at the inversion, we caught only 5 mice in this region that were homozygous for the prairie allele. To aid interpretability, we therefore coded genotype as a factor with the homozygous forest genotype as the baseline and separate effects for the heterozygote and homozygous prairie genotypes rather than separately coding the additive and dominance effects of the locus.

Absolute allele frequency differences across the genome

To determine whether the inversion frequency change across the transect was consistent with divergent selection, we compared it with blocks of similar linkage disequilibrium across the genome. We binned the genome into 200-bp non-overlapping windows, as mean $r^2$ is approximately 0.1 at this distance. Although the mean $r^2$ within the inversion is higher (~0.2), the rapid decay of linkage disequilibrium in the forest and prairie populations makes a window size of mean $r^2$=0.2 infeasible. For each 200-bp window, we considered only SNPs with MAF>0.1 and reported the SNP with highest absolute allele frequency difference between the wild-caught forest (n = 15) and prairie (n = 15) mice, with a few exceptions. In particular, we noticed three other high-LD blocks that will be subject to further investigation. We collapsed these three blocks of high-LD, as well as the inversion on chromosome 15, to single values representing the most frequent maximum absolute allele frequency difference of 200-bp windows within each block. We then compared the absolute allele frequency difference of the inversion (0.9) to the maximum absolute allele frequency difference of all similar linkage disequilibrium blocks genome-wide (Figure S11).

**Inversion frequency and associations across the species range**

Genotyping the inversion across populations

Using 3 published datasets and museum specimens, we tested for the presence/absence of the chromosome 15 inversion in 251 mice from 37 populations across North America (Baier 2019: n = 58 (*42*), Kingsley & Kozak 2017: n = 73 (*43*), Schweizer 2019: n = 96 (*44*), Burke Museum: n = 24). Data on sample number, location, genotype, dataset and sequencing type are provided in Data S3. We genotyped 24 specimens from Washington state (University of Washington, Burke Museum), using the 4 Taqman SNP genotyping assays as described above. For the published datasets, we obtained fastq files on the NCBI Sequence Read Archive (PRJNA508981 (*42*), PRJNA528923 (*44*)), or from the authors (*43*), and mapped sequencing reads to the *P. maniculatus bairdii* reference genome. We then performed the fit-HMM step of MSG (*8*) on the affected region of chromosome 15 as described above, using the set of 37,242 fixed SNPs between the inversion and reference haplotypes.

To confirm the MSG inversion genotypes, we performed a genetic principal components analysis (PCA) on SNP data from the 3 published datasets (Figure S17). Using the mapped bam files for the 227 samples, we performed joint genotyping as described above. For each dataset, we selected SNPs with at least 50% of samples genotyped, and then performed PCA using *SNPRelate* (*45*) including the forest (n = 15) and prairie (n = 15) mice. The number of SNPs used for each dataset differed by sequencing method as follows (affected; unaffected): Schweizer 2019: 31,359; 34,415; Baier 2019: 5,444; 5,982; Kingsley & Kozak 2017: 601; 515.

<u>Habitat characterization for North American populations</u>

We characterized local habitats following methods described in (*43*). For each of the 251 *P. maniculatus* mice from across North America plus the forest (n = 15) and prairie (n = 15) mice, we extracted land cover information in a 1 km radius around the reported capture site from the North American Land Change Monitoring System (NALCMS 2010) with 30m spatial resolution. We then recorded the percent of pixels (*percent-forested*) in the 1 km radius that were categorized as forest habitat (habitat class in mixed_forest, temp_subpolar_needleleaf_forest, temperate_subpolar_broadleaf_deciduous_forest, subpolar_taiga_needleleaf_forest, tropical_subtropical_broadleaf_deciduous_forest, or tropical_subtropical_broadleaf_evergreen_forest).

<u>Inversion associations with habitat and tail length</u>

To test for associations between the inversion and tail length or habitat across the species range, we implemented two approaches to account for population structure in the set of genotyped mice: (1) using a kinship matrix as a random effect, and (2) using genetic principal components as covariates in linear models.

We built a kinship matrix with EMMAX (v.07Mar2010) (*46*) using SNP data. From the vcf from across all datasets with *P. maniculatus* mice, we selected SNPs with > 80% of samples genotyped, and we excluded Baier 2019 mice so that we had a sufficient number of overlapping SNPs across datasets. We converted the vcf to a bed file and then a tped file using *plink,* and then built an IBS kinship matrix using EMMAX. For the kinship matrix, we used a total of 3,427 SNPs across all autosomes except chromosome 15. We excluded chromosome 15 so that the inversion genotypes did not affect kinship coefficients. We then tested for an association between inversion genotype and habitat (*percent-forested*) using EMMAX, with inversion genotype as the only input genotype, the kinship matrix as a random effect and habitat as the response variable. We also tested for an association between inversion genotype and tail length. We obtained tail length data for a total of 111 mice: 29 from prairie and forest populations, 67 from the Kingsley & Kozak 2017 dataset from the author, and 15 mice from the Schweizer 2019 dataset that had museum accession numbers. We implemented the linear mixed model using EMMAX, with tail length as the response variable, inversion genotype as the only input genotype, body length as a fixed effect and the kinship matrix as a random effect.

As a second approach for the association analysis, we performed a genetic PCA using *SNPRelate* (*45*) to account for population structure. For PCA, we used the same set of SNPs used for the kinship matrix (3,427 SNPs across all autosomes except chromosome 15), and we note that the top PCs separated mice by population (Figure S18). We then implemented linear models in R with inversion genotype as a fixed numeric effect and the top 5 principal components as fixed effects. Habitat (*percent-forested)* and tail length were the response variables, and for tail length, we included body length as an additional fixed effect. For both habitat and tail length, we performed a likelihood ratio test between models with and without inversion genotype using the *anova* function in R (tail length: $p = 9.7e-8$; habitat: $p < 2.2e-16$).

**Permissions and approval**

We trapped mice under Oregon Department of Fish and Wildlife Scientific Taking Permits #107-15 and 127-16, with approval from Siuslaw, Deschutes, and Willamette National Forests and the Bureau of Land Management. We imported colony founders to Massachusetts on Division of Fisheries and Wildlife Importation Permit #043.15IMP. All experiments were approved by Harvard's IACUC (Protocol 11-05).

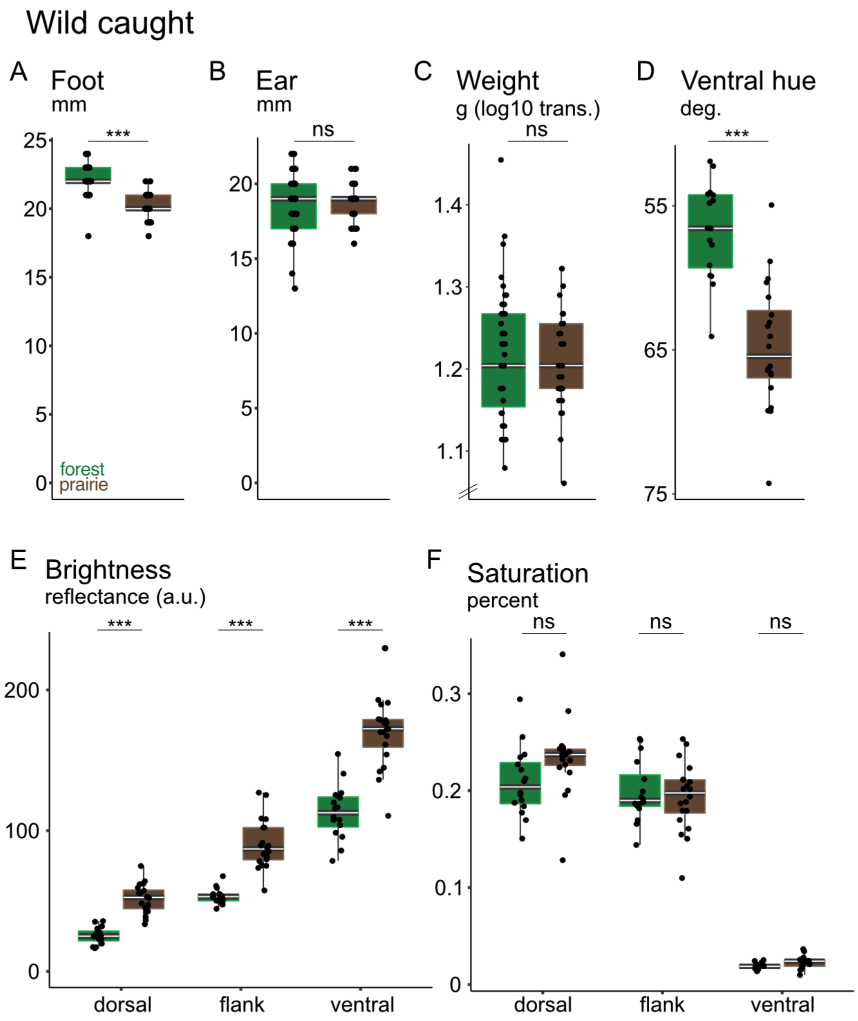

**Fig. S1.**

**Additional phenotypes of wild-caught mice.** Phenotypes of adult, wild-caught mice from the forest (green, left) and prairie (brown, right) ecotypes. (A) Hindfoot length (n = 33 forest, 29 prairie), (B) ear length (n = 33 forest, 29 prairie), (C) weight (shown after $\log_{10}$ transformation; n = 39 forest, 30 prairie), and (D) ventral hue. (E) brightness and (F) saturation of all three body regions (n = 16 forest, 20 prairie for all pigment traits). Symbols: ns = $p > 0.05$; *** = $p < 0.001$ (two-sided Welch's t-tests); deg. = degrees; a.u. = arbitrary units of reflectance.

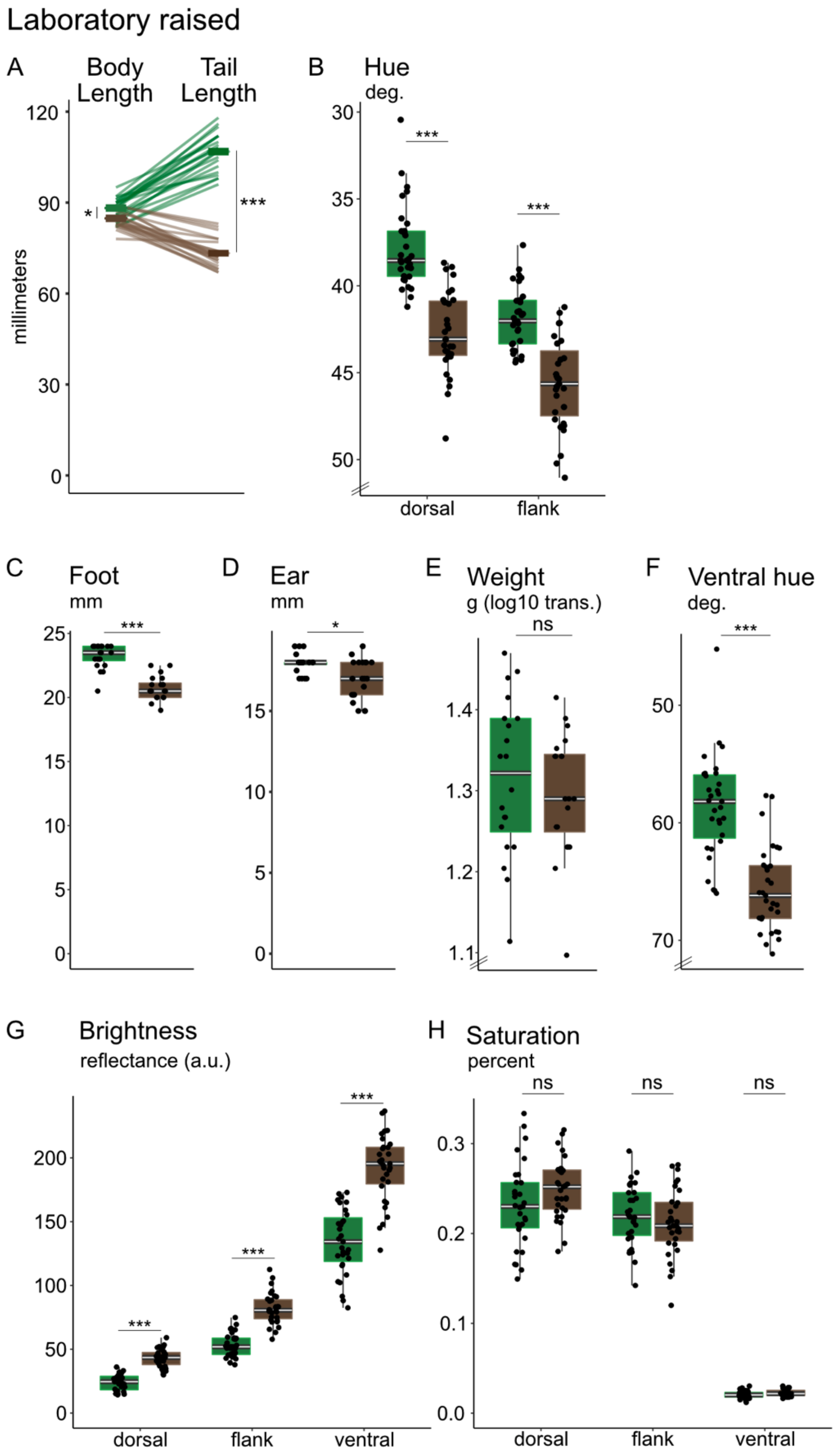
Laboratory raised
A
Body Length
Tail Length
millimeters
B
Hue
deg.
dorsal
flank
C
Foot
mm
D
Ear
mm
E
Weight
g (log10 trans.)
ns
F
Ventral hue
deg.
G
Brightness
reflectance (a.u.)
dorsal
flank
ventral
H
Saturation
percent
ns
dorsal
flank
ventral

**Fig. S2.**

**Phenotypes of laboratory-born mice are consistent with wild-caught specimens.** Phenotypes of laboratory-born adult forest (green) and prairie (brown) mice, aged 60-70 days (n = 20 forest, 20 prairie for body measurements, n = 31 forest, 31 prairie for pigment traits). (A) Body length, excluding the tail, and tail length. Lines connect measurements for the same individual. (B) Dorsal and flank hue. (C) Hindfoot length, (D) ear length, (E) weight (after $\log_{10}$ transformation), and (F) ventral hue. (G) Brightness and (H) saturation for all three body regions. Symbols: ns = $p > 0.05$; * = $p < 0.05$; *** = $p < 0.001$ (two-sided Welch's t-tests); deg. = degrees; a.u. = arbitrary units of reflectance.

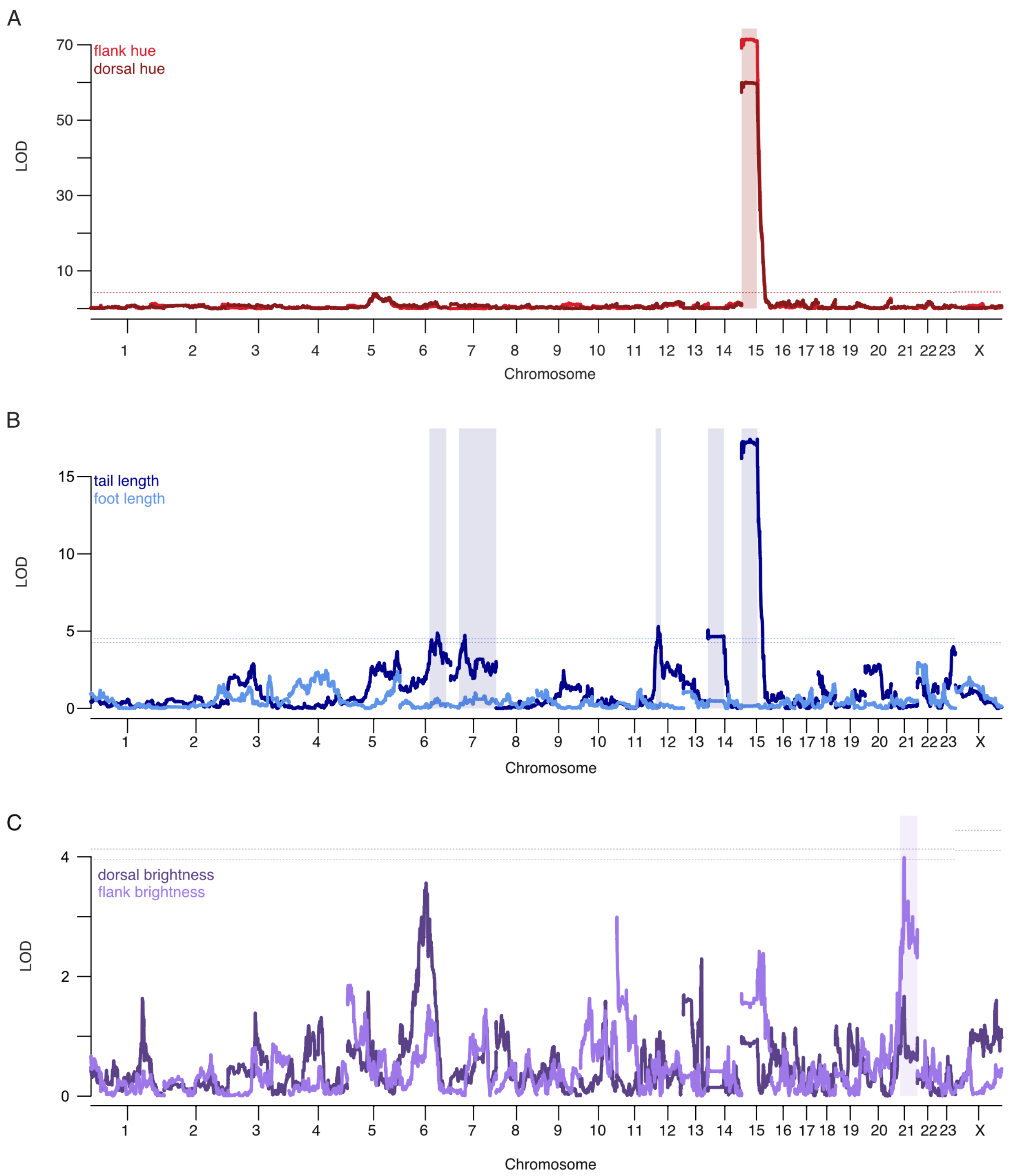

**Fig. S3.**

**QTL maps for all traits that differed between wild-caught forest and prairie mice.** (A) Flank hue (light red) and dorsal hue (dark red). (B) Tail length (dark blue) and hindfoot length (light blue). For tail and hindfoot length analyses, body length was included as an additive covariate. (C) Dorsal brightness (dark purple) and flank brightness (light purple). The peak for flank brightness on chromosome 21 is transgressive. LOD = log of the odds score. Physical distance in basepairs is shown on the x-axis; axis labels indicate the center of each chromosome. Dotted lines indicate the genome-wide significance threshold ($\alpha = 0.05$) based on permutations, and shaded rectangles indicate the 95% Bayes' credible intervals for all chromosomes with significant QTL peaks. n = 542 (tail), 455 (foot), 541 (pigment).

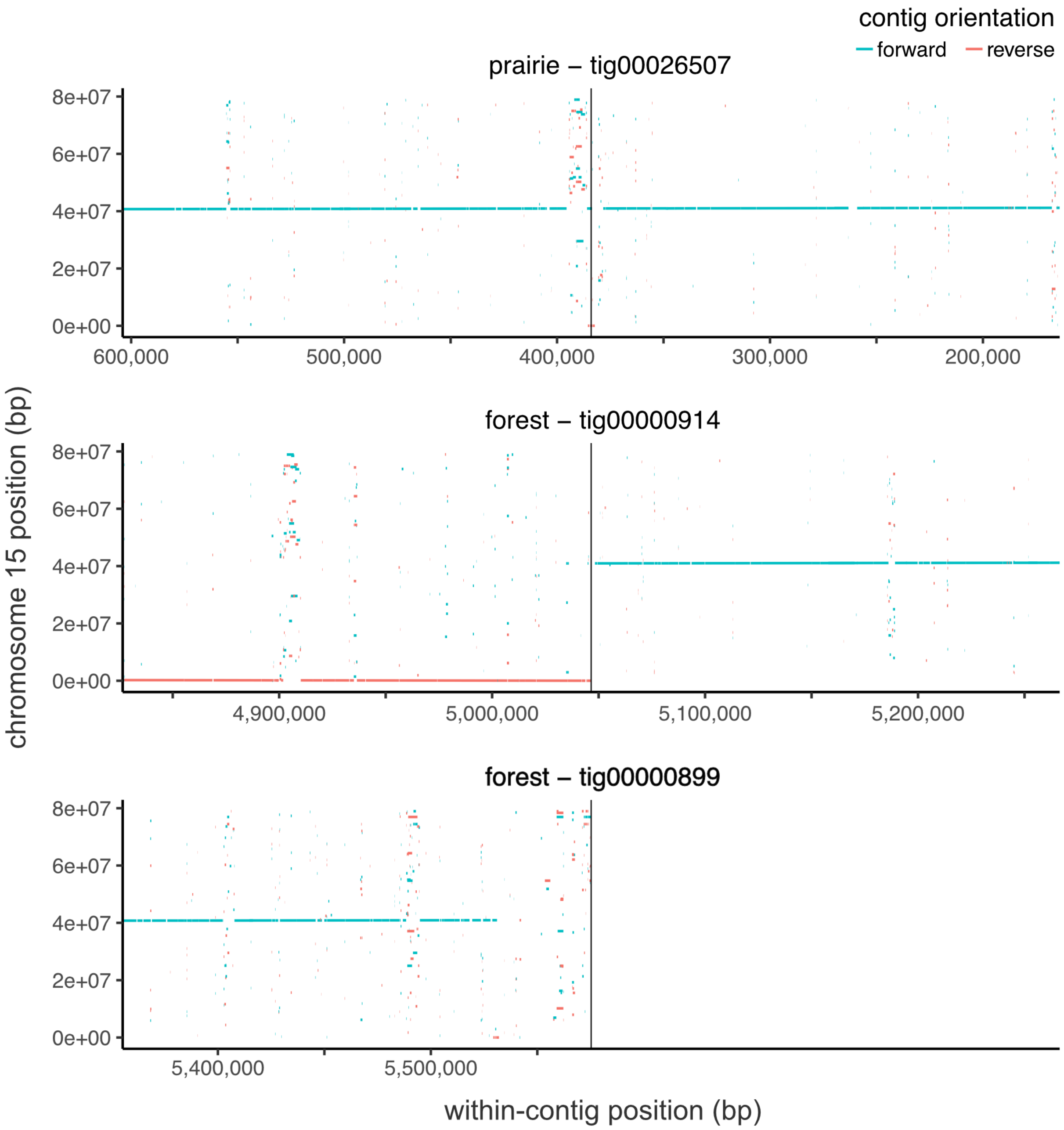

**Fig. S4.**

**Contig alignments: repetitive region**. Alignments of the three long-read sequencing-based contigs relevant for the inversion breakpoint: the prairie contig spanning the breakpoint (top), the forest contig spanning the breakpoint (middle), and the forest contig that contains sequence that is adjacent to the breakpoint (bottom). Contigs are aligned to chromosome 15 from the *P. maniculatus* reference genome (y-axis). Vertical line: identified breakpoint. Colors indicate alignment direction with respect to the *P. maniculatus* reference genome.

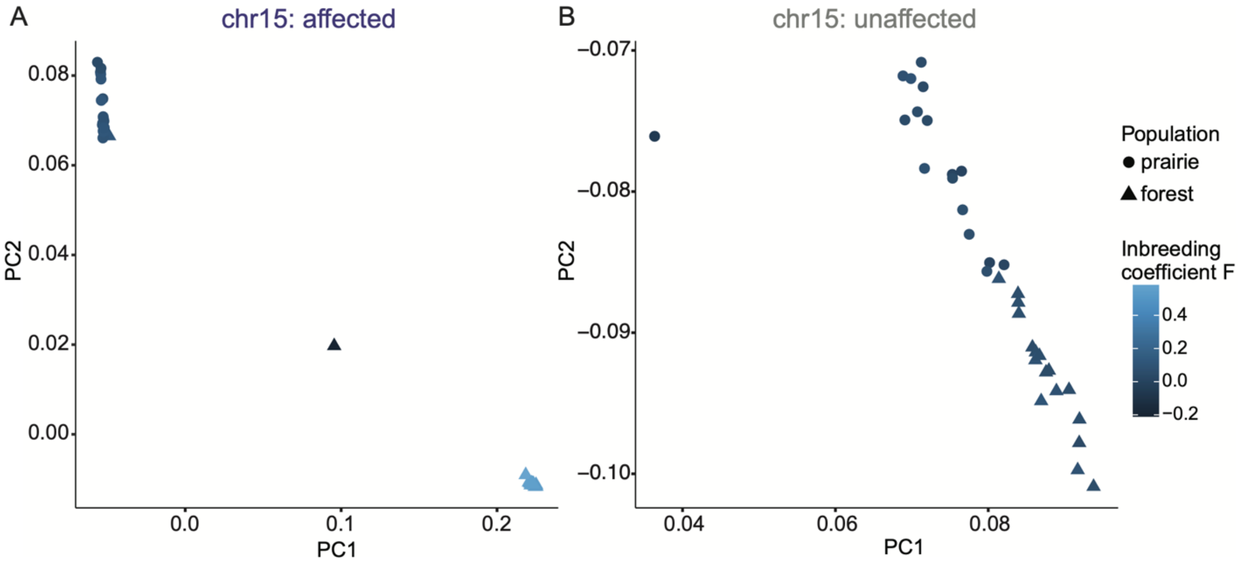

**Fig. S5.**

**Genetic principal component analyses and heterozygosity of the affected and unaffected regions of chr15.** Genetic PCA performed for (A) the affected and (B) the unaffected regions of chromosome 15, using wild caught prairie and forest populations. Each point represents an individual and is colored by the inbreeding coefficient F. The structure of the affected region portions out individuals into three distinct clusters, which is congruent with heterozygosity and inversion genotypes, representing homozygous reference individuals (left), a single heterozygous individual (middle), and homozygous inversion individuals (right). The unaffected region shows little variation in F, and structures samples primarily by source population (forest v. prairie).

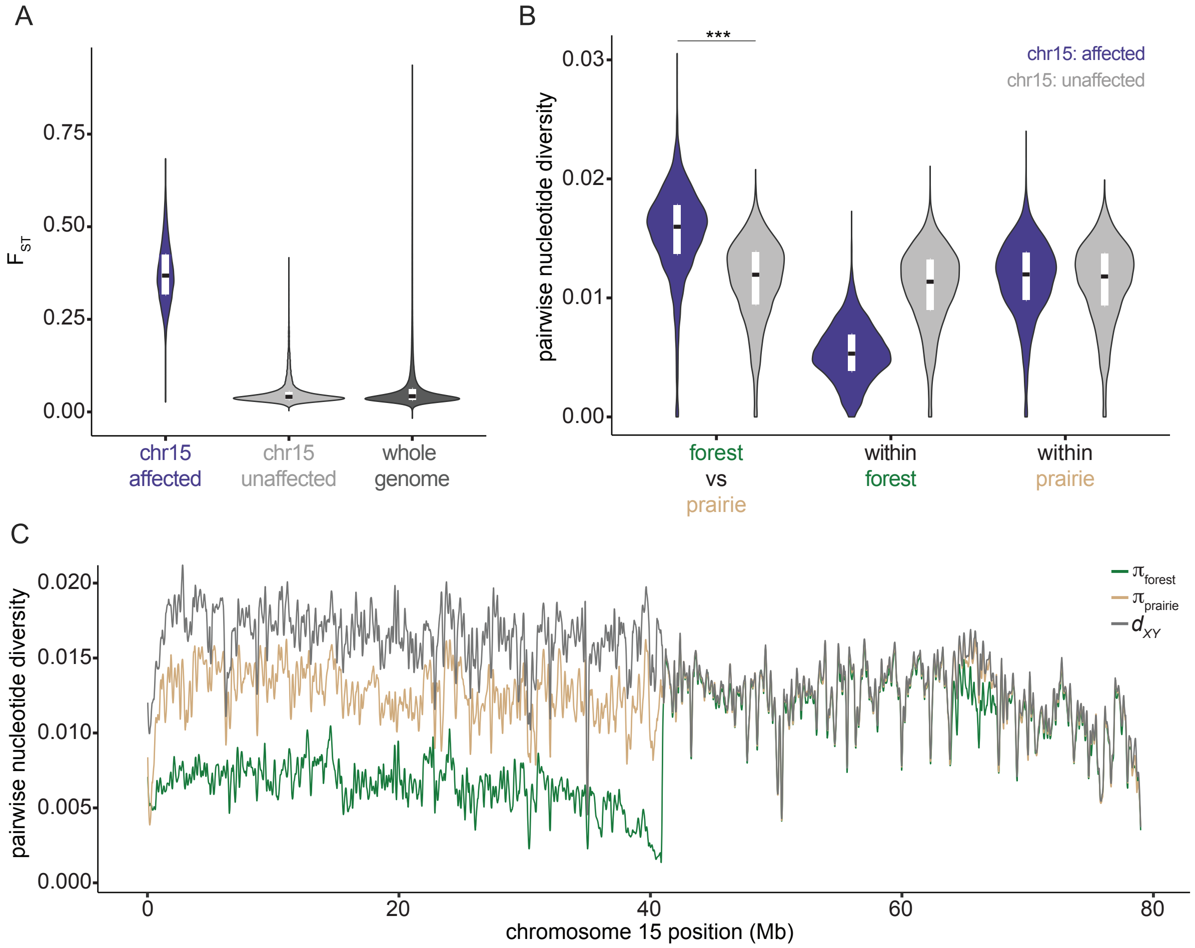

**Fig. S6.**
**Genetic differentiation statistics for chromosome 15.** (A) $F_{ST}$ between wild-caught forest (n = 15) and prairie (n = 15) mice calculated in 10-kb windows with step size of 1 kb in the affected (purple, chr15:0-40 Mb) and unaffected (light gray, chr15:41-79 Mb) regions of chromosome 15, and across the whole genome excluding the affected region of chromosome 15 (dark gray). (B) Pairwise nucleotide diversity for forest v. prairie ecotypes ($d_{XY}$), within forest mice ($\pi_{forest}$), and within prairie mice ($\pi_{prairie}$) shown for the affected (purple, chr15:0-41 Mb) and unaffected (light gray, chr15:41-79 Mb) regions of chromosome 15. Nucleotide diversity statistics were computed in 10-kb windows with step size of 10 kb. Only forest mice homozygous for the inversion were included in nucleotide diversity analyses. For all violin plots, white boxes represent first and third quartiles, with median shown as black line. Symbols: *** = $p < 0.001$ (two-sided Student's t-test) (C) Smoothed nucleotide diversity shown across chromosome 15 (green = $\pi_{forest}$, tan = $\pi_{prairie}$, gray = $d_{XY}$).

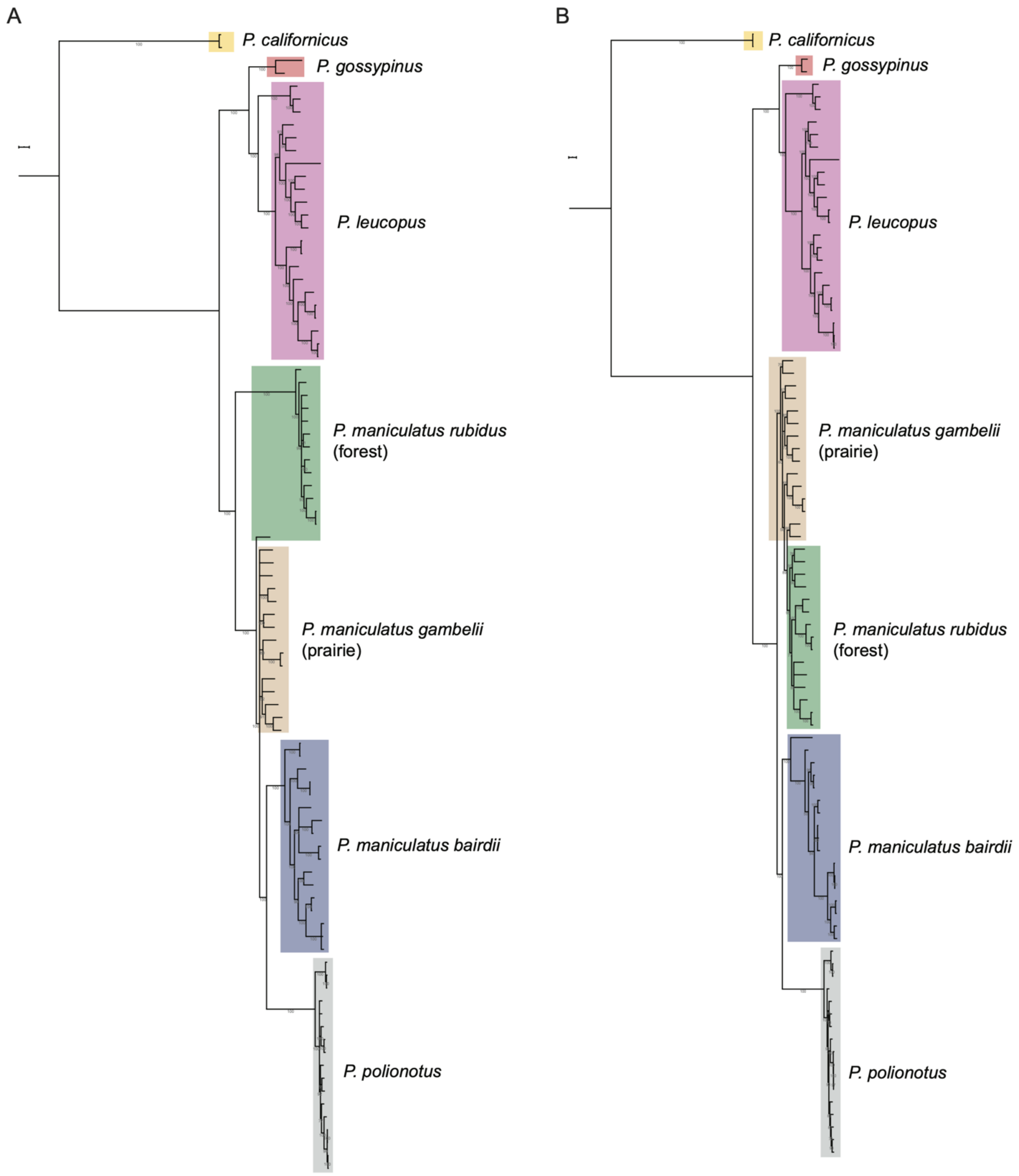
A
P. californicus
P. gossypinus
P. leucopus
P. maniculatus rubidus
(forest)
P. maniculatus gambelii
(prairie)
P. maniculatus bairdii
P. polionotus
B
P. californicus
P. gossypinus
P. leucopus
P. maniculatus gambelii
(prairie)
P. maniculatus rubidus
(forest)
P. maniculatus bairdii
P. polionotus

**Fig. S7.**

**Maximum likelihood trees for *Peromyscus* species.** Maximum likelihood trees of *Peromyscus* species showing all mice for affected (A) (chr15:0-40.9 Mb) and unaffected (B) (chr15:40.9-79 Mb) regions of chromosome 15. Branches with < 50 bootstrap support are collapsed. Shaded boxes indicate mice belonging to a single species or subspecies as labeled.

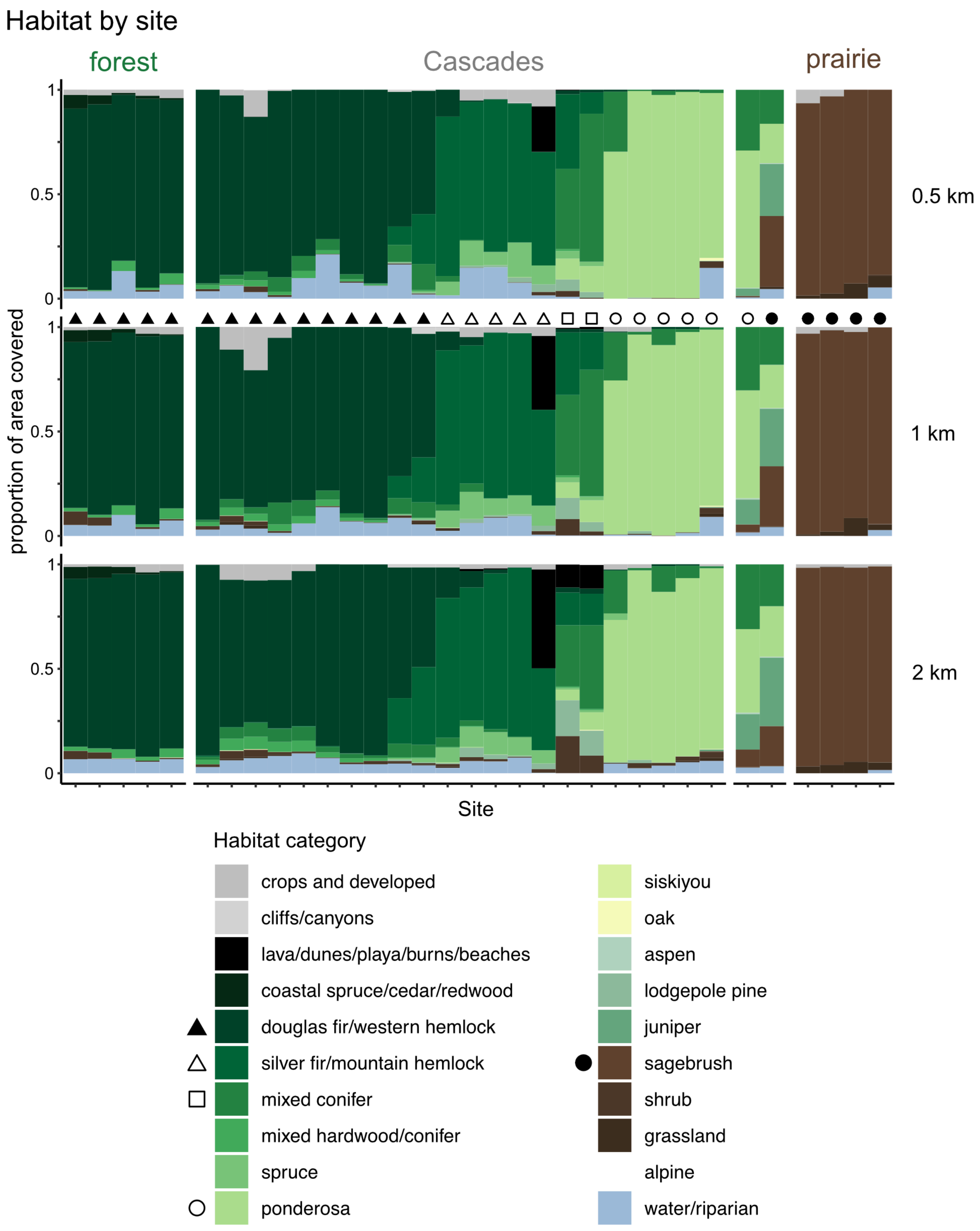

Habitat by site
forest
Cascades
prairie
proportion of area covered
0
0.5
1
0.5 km
1 km
2 km
Site
Habitat category
crops and developed
cliffs/canyons
lava/dunes/playa/burns/beaches
coastal spruce/cedar/redwood
douglas fir/western hemlock
silver fir/mountain hemlock
mixed conifer
mixed hardwood/conifer
spruce
ponderosa
siskiyou
oak
aspen
lodgepole pine
juniper
sagebrush
shrub
grassland
alpine
water/riparian

**Fig. S8.**

**Habitat cover at sampled sites.** The proportion of the area within 0.5 km (top), 1 km (middle) and 2 km (bottom) covered by each habitat category. Data was drawn from the habitat map by the Oregon Biodiversity Information Center of the Institute for Natural Resources at Portland State University, and habitat categories were binned across age categories as shown in Table S4. The habitat categories shown in Figure 5 represent the habitat that covers the most area within 1 km of the site; symbols shown correspond to those in Figure 5.

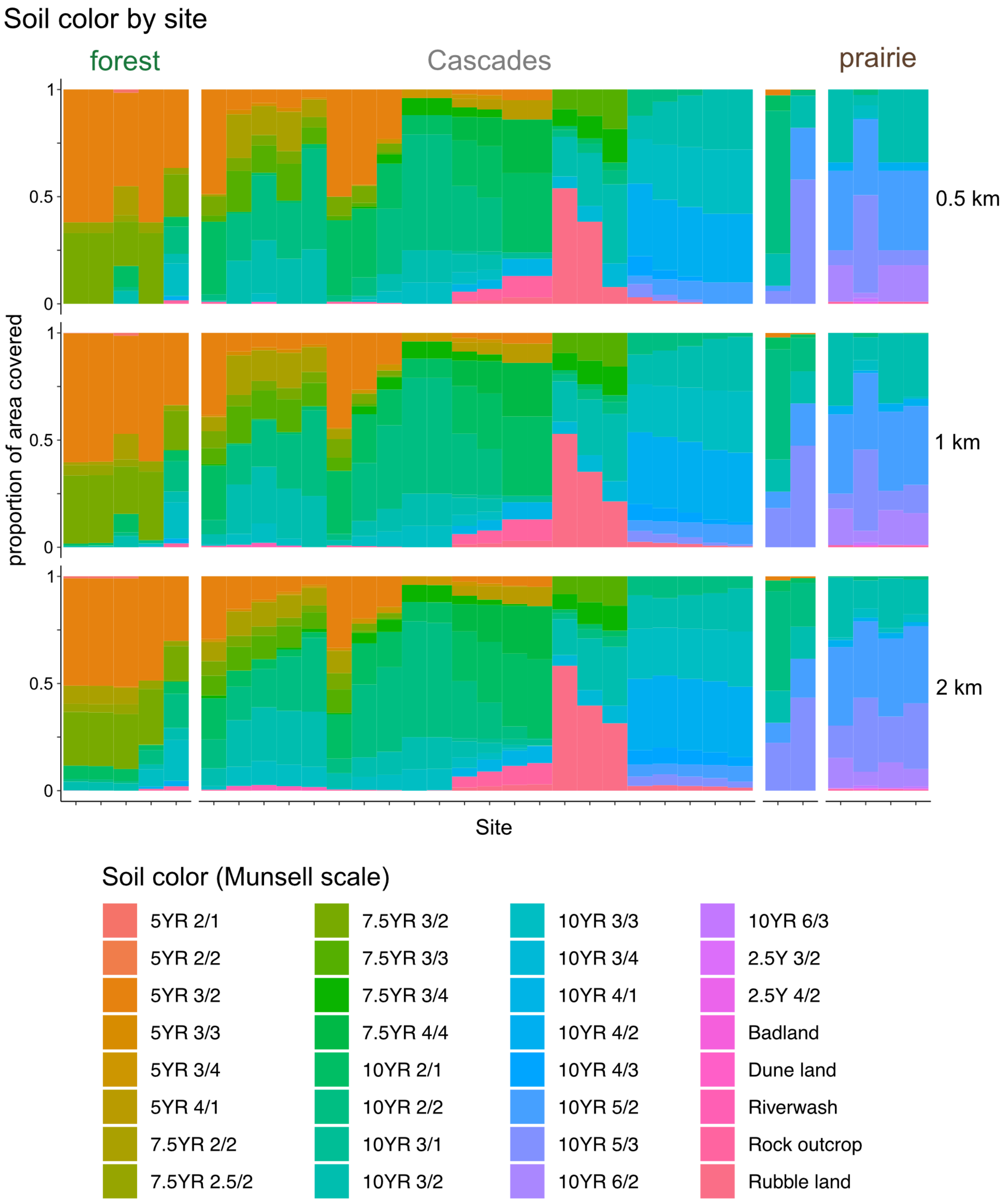

**Fig. S9.**

**Soil characteristics at sampled sites.** The proportion of top-layer soil with each Munsell scale color (shown as hue value/chroma), within 0.5 km (top), 1 km (middle), or

2 km (bottom) of each sampled site. The sites are ordered by transect distance (i.e., distance east from the central point). The hue values shown in Figure 5 are the weighted average of the Munsell hue for the 1 km radius shown here, after excluding regions with no soil series data (i.e. badland, dune land, riverwash, rock outcrop, and rubble land categories).

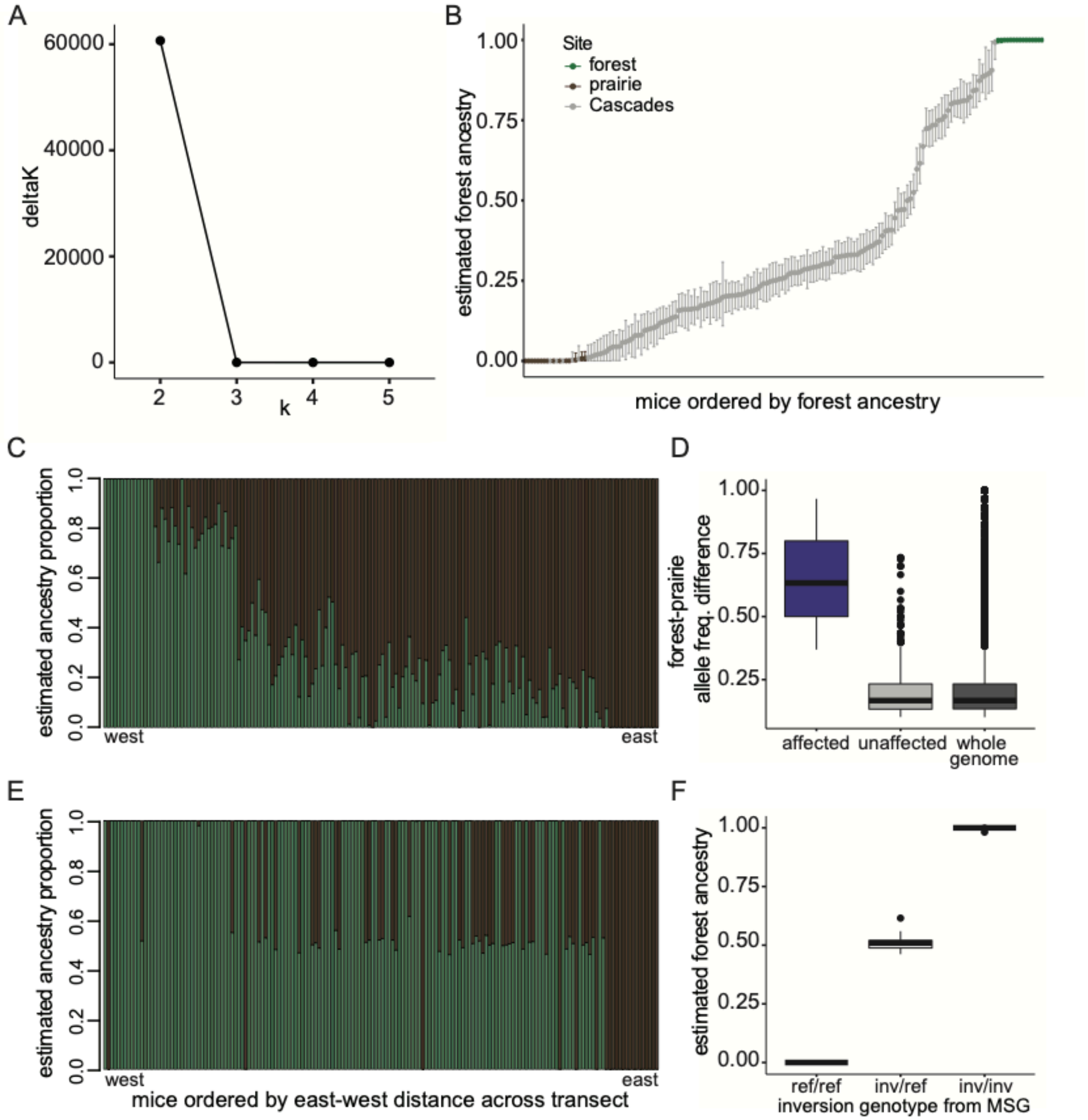

**Fig. S10.**

**ngsAdmix ancestry estimates.** (A) Results from the Best K method (CLUMPAK) shows that ancestry for all transect mice was best assigned to two clusters (k=2). (B) Bootstrapped confidence intervals for genome-wide ngsAdmix ancestry estimates. Points show percent of ancestry assigned to cluster 1 (forest ancestry) with the full SNP set, and bars show 95% confidence interval on forest ancestry from bootstrapping. Mice are ordered by their forest ancestry estimates from full SNP set, and colored by site (forest=green, prairie=brown, Cascades=gray). (C) Genome-wide ngsAdmix ancestry estimates for forest, Cascades and prairie mice for all autosomes, excluding affected region of chromosome 15. Mice are ordered by their distance along an east-west axis of their capture sites, with western sites on the left and eastern sites on the right. All forest mice (n = 15, left) are assigned 100% ancestry in cluster 1 whereas all prairie mice (n = 15, right) are assigned 100% ancestry in cluster 2, suggesting that the two ancestry groups correspond to forest ancestry (green) and prairie ancestry (brown). Central Cascades transect mice have varying proportions of ancestry assignments (middle). (D) Allele frequency difference between forest and prairie populations for the top quartile of differentiated SNPs that were used in ngsAdmix for determining genome-wide ancestry (whole genome) and ancestry for the inversion region (affected). ngsAdmix SNPs from the unaffected region of chromosome 15 (unaffected) are also shown for comparison with the affected region. (E) ngsAdmix ancestry estimates for the affected region of chromosome 15 to confirm inversion genotypes, with mice ordered as in (C). (F) ngsAdmix estimates for forest ancestry at the affected region agree with inversion genotypes as determined by MSG.

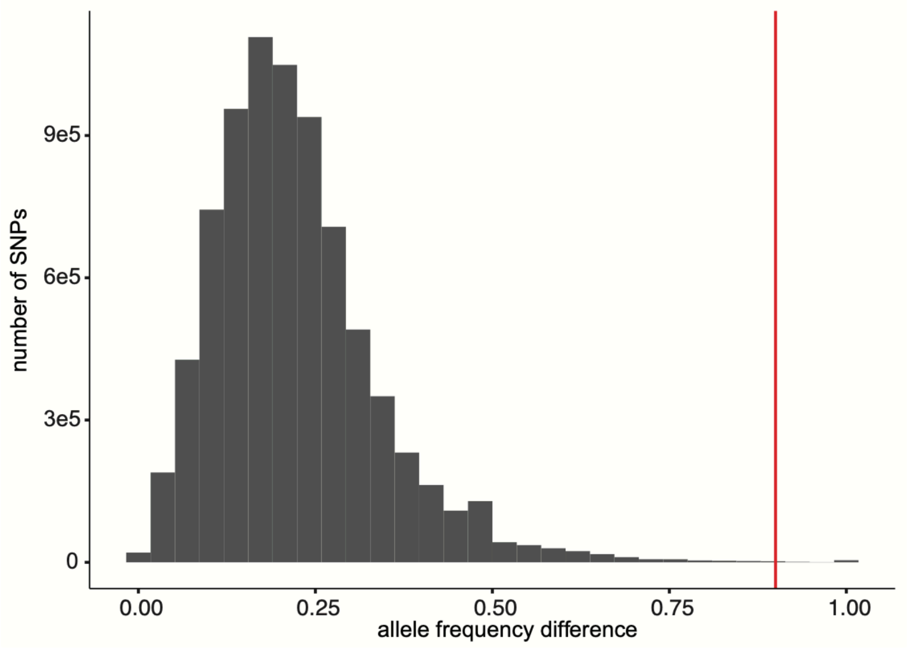

**Fig. S11.**
**Absolute forest-prairie allele frequency difference for sites genome-wide.** Allele frequency differences for the maximally differentiated SNP between forest and prairie mice in 200-bp windows across the genome. The inversion allele frequency difference between forest and prairie mice of 90% is shown in red.

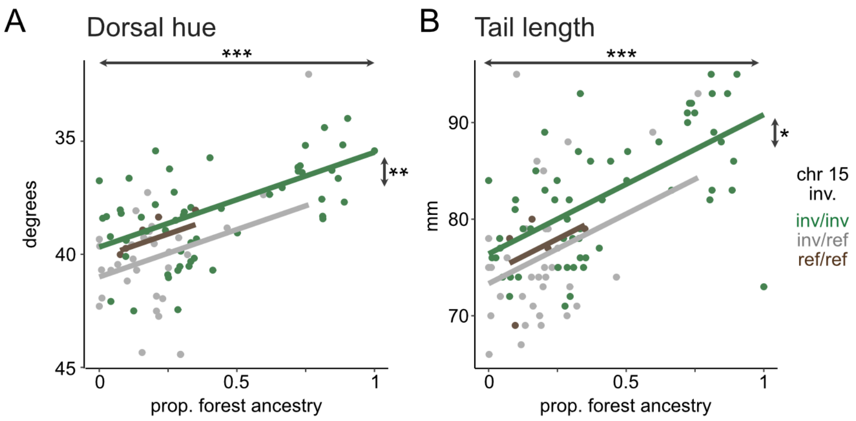

**Fig. S12.**

**Association between genotype and phenotype in transect mice.** Relationship between estimated forest ancestry proportion (x axis), genotype at the chromosome 15 inversion (green = homozygous for the inverted (forest) allele; gray = heterozygous; brown = homozygous for the reference (prairie) allele), and dorsal hue (A) and tail length (B) in wild-caught adults from the central Cascades transect. Points show individual mice (dorsal hue: n = 90; tail: n = 97), and lines show the results of mixed-effect models including capture site as a random effect. In both cases, models including both inversion genotype and genome-wide ancestry were selected over equivalent models with genome-wide ancestry only (tail, inversion + ancestry, model log-likelihood (LL) = -304.4, AIC = 620.9; tail, ancestry only, LL = -307.0, AIC = 622.0; hue, inversion + ancestry, model log-likelihood (LL) = -174.6, AIC = 361.3; hue, ancestry only, LL = -179.0, AIC = 366.0). Symbols: * = $p < 0.05$; ** = $p < 0.01$; *** = $p < 0.001$.

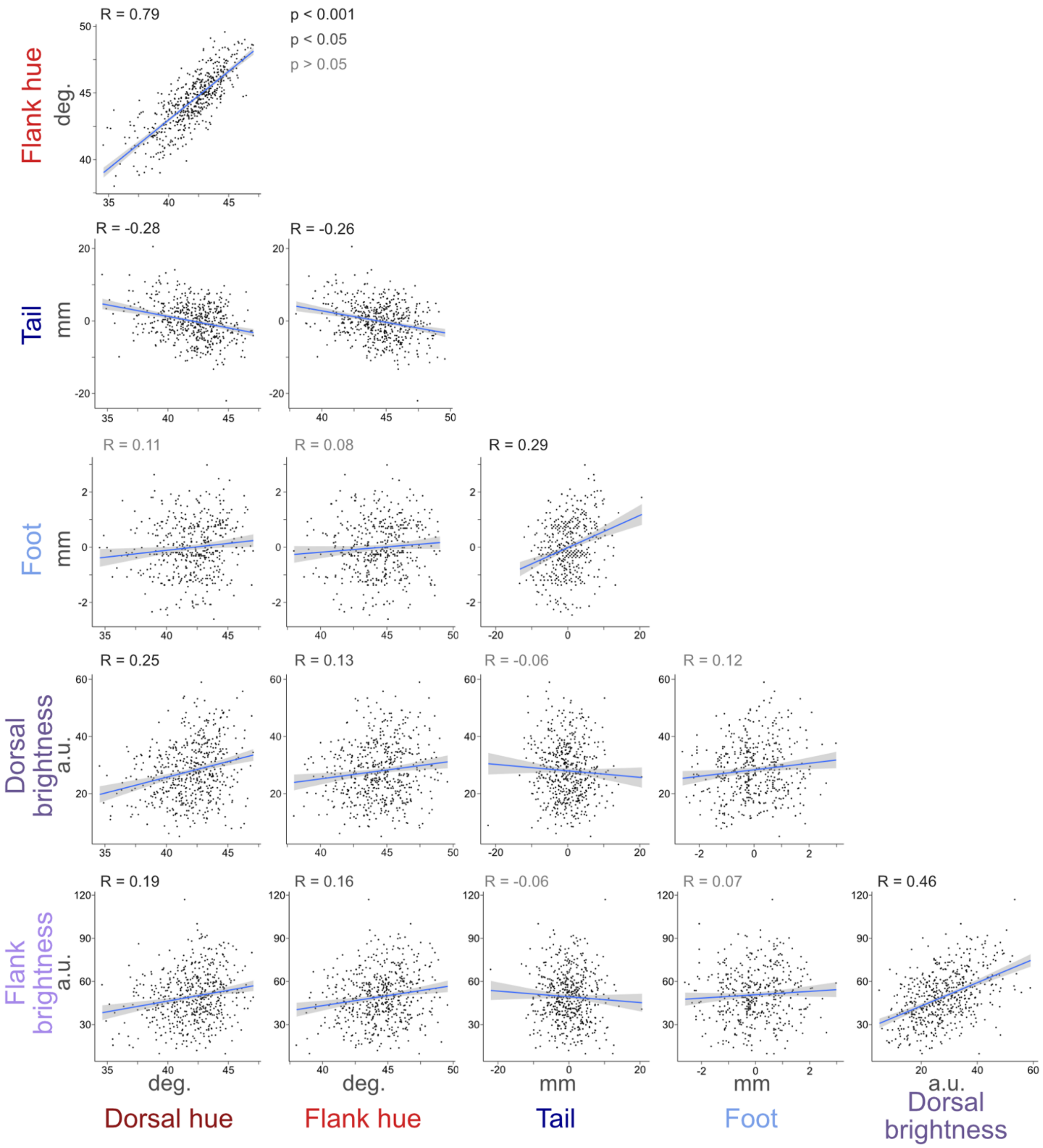

**Fig. S13.**

**Correlations among traits in F2 hybrids.** Pairwise Pearson's correlations among all traits used for QTL mapping in F2 hybrids. Tail and hindfoot length are shown after taking the residual against body length in the hybrid mice. n = 542 (tail), 455 (foot), 541 (pigment).

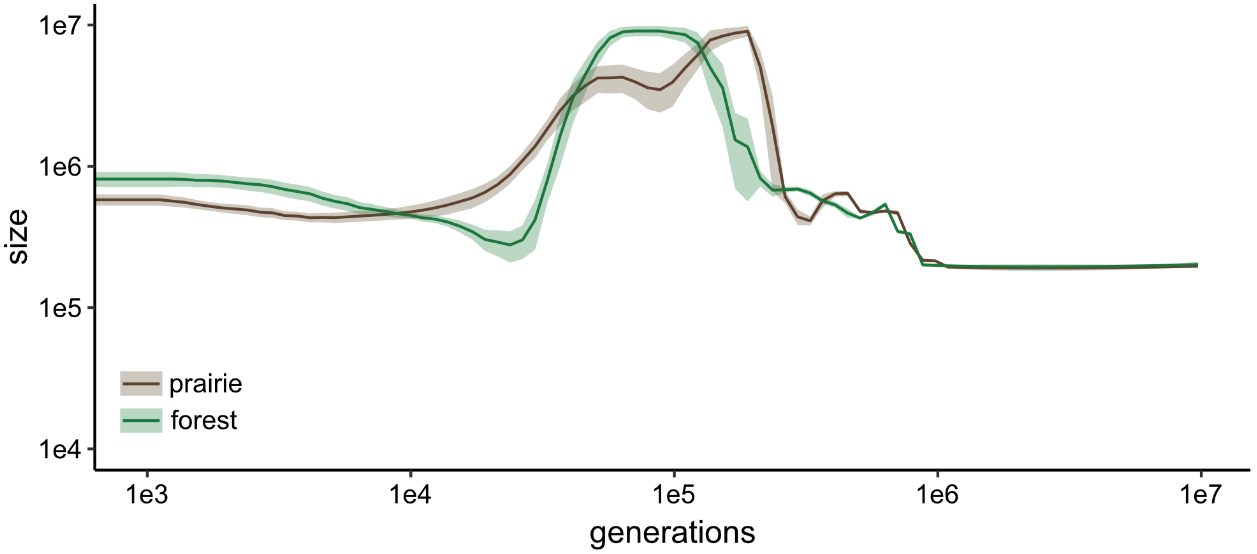

**Fig. S14.**

**Demographic history through time of prairie and forest populations.** Log-scaled population size trajectories through time, with most recent history plotted on the left. Solid lines represent mean sizes and shaded ribbons 95% CI calculated from original data and 20 bootstrap replicates for each population.

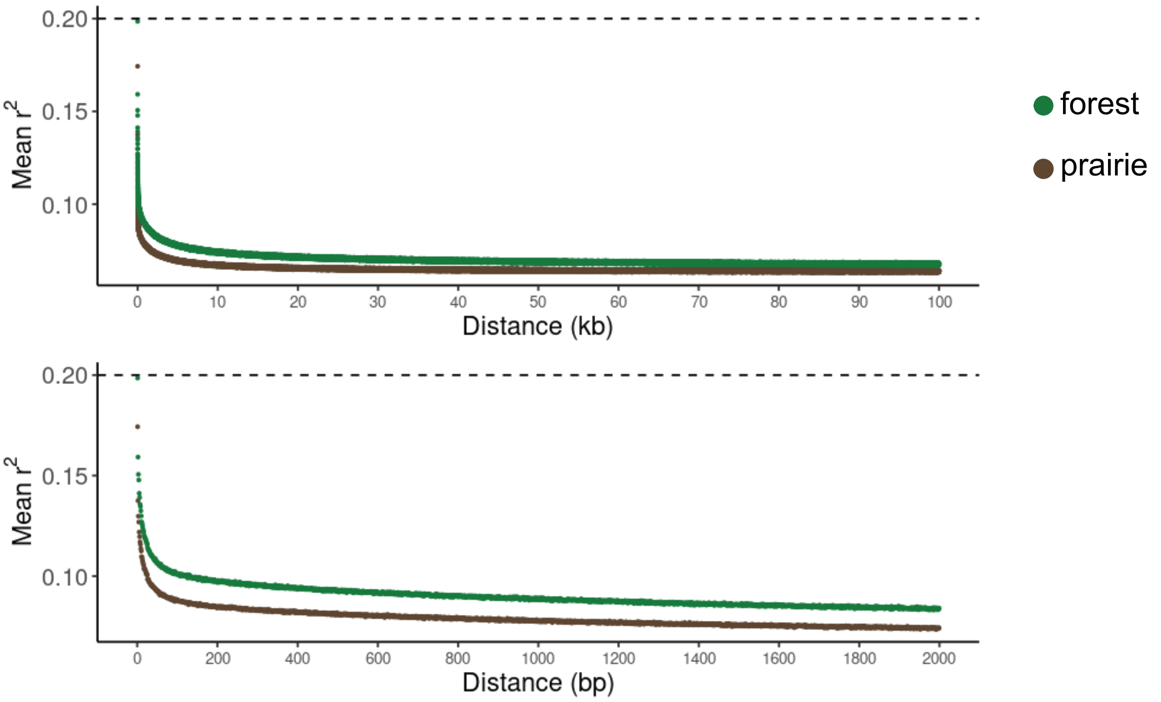

**Fig. S15.**

**LD decay in wild prairie and forest populations.** Mean $r^2$ between biallelic SNPs, as a function of physical distance in kb (top) and base pairs (bottom). Horizontal line at $r^2$ = 0.20 indicates a common threshold below which LD between variants is often considered to be negligible.

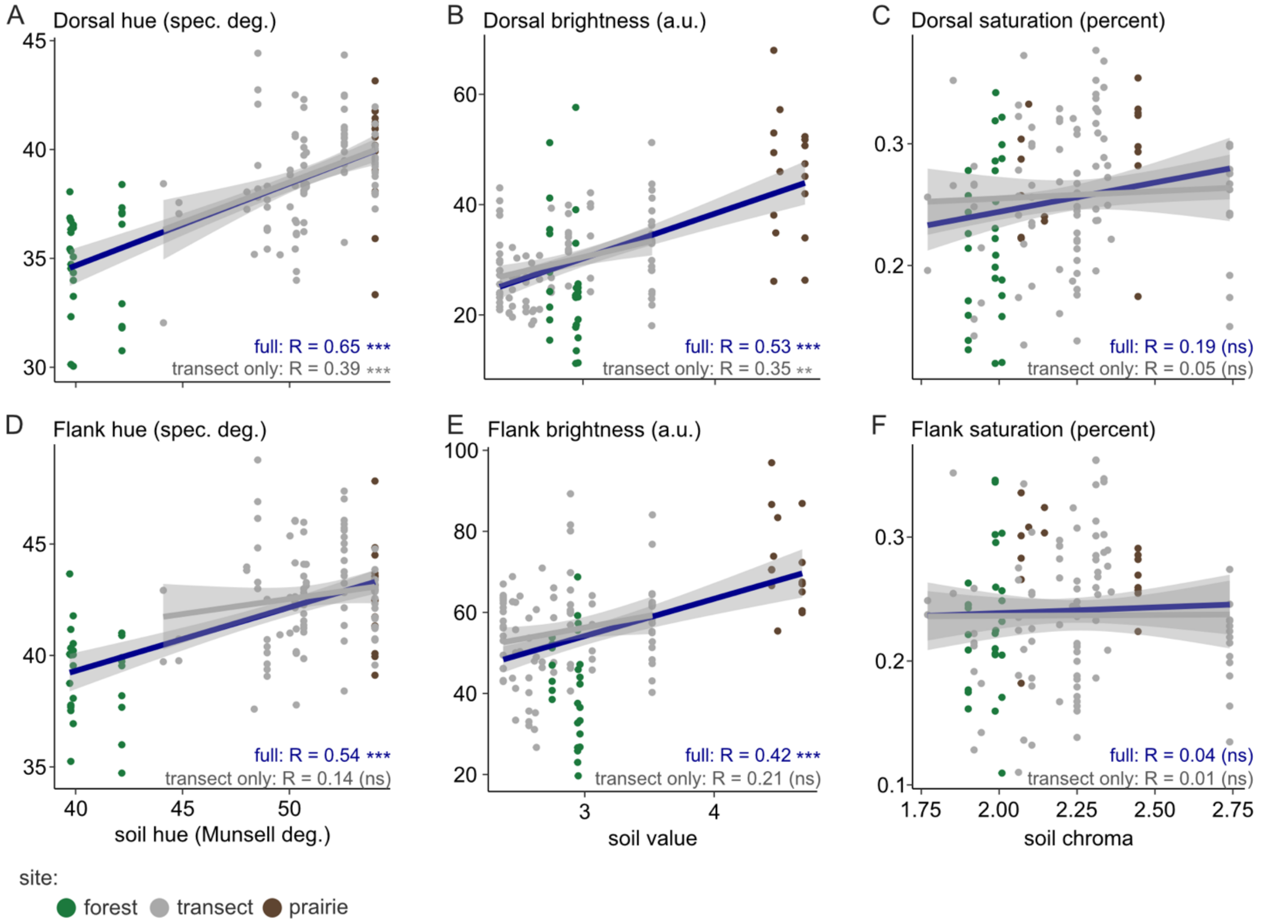

**Fig. S16.**

**Correlation between mouse pigmentation and soil characteristics.** Correlations between mean soil hue (A,D), value (B,E), and chroma (C,F) and dorsal (A-C) or flank (D-F) hue, brightness, and saturation in the wild-caught mice. Points are colored by location of capture (green = forest, gray = transect, brown = prairie). Correlations are shown both using all data (full = blue, n = 133) and using the central Cascades transect only (gray, n = 90). Symbols: ns = p > 0.05; ** = p < 0.01; *** = p < 0.001.

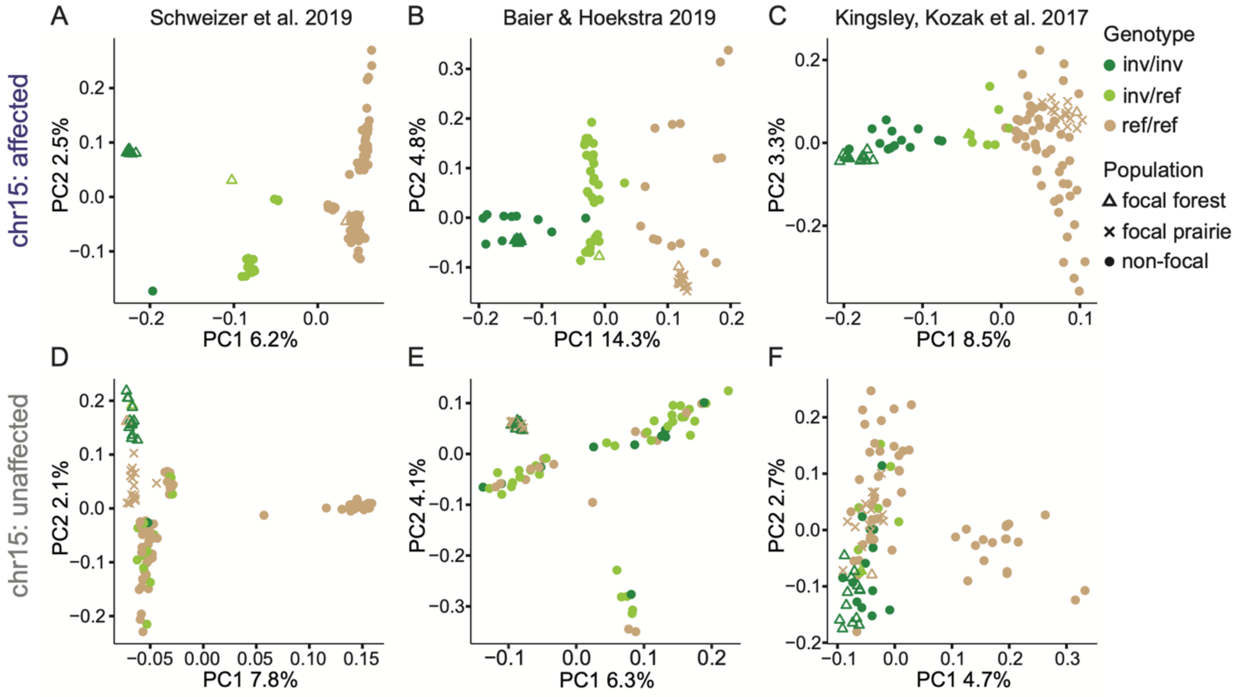

**Fig. S17.**

**Genetic principal component analyses for *P. maniculatus.*** Genetic PCA performed for chromosome 15 for *P. maniculatus* mice from the three published datasets. Axes indicate percent variance explained by top two principal components, points are colored by genotype at the inversion locus as determined by MSG, and shapes indicate the population (non-focal = samples from three published datasets). PCA plots using SNPs from the affected region of chromosome 15 are shown for the Schweizer (A), Baier (B) and Kingsley & Kozak (C) datasets. In all three datasets, PC1 separates mice by genotype at the inversion locus. PCA plots using SNPs from the unaffected region of chromosome 15 are shown for the Schweizer (D), Baier (E) and Kingsley & Kozak (F) datasets. The top two principal components do not separate mice by inversion genotype.

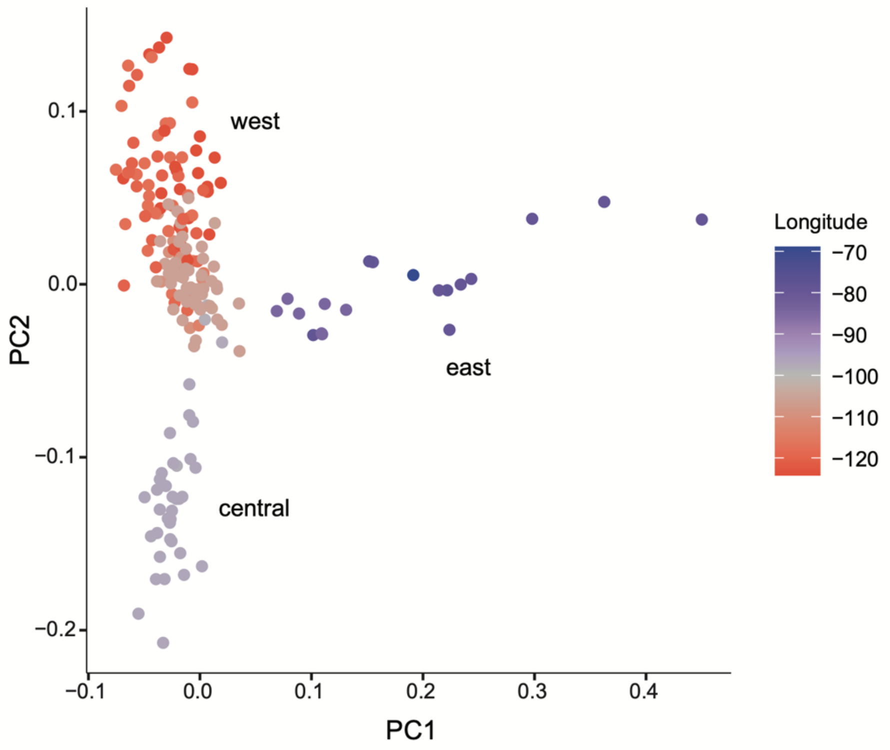

**Fig. S18.**

**Genetic PCA for *P. maniculatus* dataset.** Genetic PCA performed on focal and non-focal (Schweizer 2019, Kingsley & Kozak 2017) *P. maniculatus* mice using genome-wide SNPs, excluding chromosome 15. PC1 separates eastern mice and PC2 separates central mice from the western mice.

**Table S1.**

**Differences in phenotype between forest and prairie mice in the wild and the lab.** Results of Welch's t-tests comparing forest and prairie phenotypes. Weight: $\log_{10}$-transformed values. s.d. = standard deviation; p (adjusted) = p-value after multiple test correction using the Bonferroni-Holm method; p (initial) = p-value without multiple test correction. Symbols: mm = millimeters, g = grams, a.u. = arbitrary units of reflectance, d = dorsal, f = flank, v = ventral, ns = p > 0.05, * = p < 0.05, *** = p < 0.001 after multiple test correction.

**WILD**

| trait | N (forest) | N (prairie) | forest (mean ± s.d.) | prairie (mean ± s.d.) | t statistic | | p (adjusted) | p (initial) |
|---|---|---|---|---|---|---|---|---|
| body (mm) | 38 | 32 | 81.5 ± 7.5 | 82.9 ± 5.7 | 0.92 | ns | 1 | 0.36 |
| weight (g) | 39 | 30 | 1.22 ± 0.08 | 1.21 ± 0.06 | -0.3 | ns | 1 | 0.77 |
| ear (mm) | 33 | 29 | 18.6 ± 2.1 | 18.7 ± 1.3 | 0.26 | ns | 1 | 0.8 |
| tail (mm) | 39 | 32 | 97.2 ± 7.4 | 69.9 ± 4.6 | -19.01 | *** | 4.63E-27 | 3.31E-28 |
| hindfoot (mm) | 33 | 29 | 22.2 ± 1.2 | 20.2 ± 1.0 | -6.95 | *** | 3.37E-08 | 3.07E-09 |
| d. brightness (a.u.) | 16 | 20 | 25.3 ± 6.1 | 51.6 ± 10.4 | 9.51 | *** | 1.20E-09 | 9.26E-11 |
| f. brightness (a.u.) | 16 | 20 | 53.4 ± 5.7 | 90.4 ± 17.4 | 8.92 | *** | 4.68E-08 | 4.68E-09 |
| v. brightness (a.u.) | 16 | 20 | 113.2 ± 19.3 | 169.7 ± 24.8 | 7.69 | *** | 5.46E-08 | 6.07E-09 |
| d. hue (degrees) | 16 | 20 | 40.3 ± 1.9 | 45.7 ± 2.0 | 8.38 | *** | 1.23E-08 | 1.03E-09 |
| f. hue (degrees) | 16 | 20 | 44.7 ± 2 | 48.9 ± 2.2 | 5.95 | *** | 7.31E-06 | 1.04E-06 |
| v. hue (degrees) | 16 | 20 | 56.7 ± 3.3 | 64.7 ± 4.4 | 6.24 | *** | 3.45E-06 | 4.31E-07 |
| d. saturation (%) | 16 | 20 | 0.21 ± 0.04 | 0.23 ± 0.04 | 2.03 | ns | 0.25 | 0.05 |
| f. saturation (%) | 16 | 20 | 0.20 ± 0.03 | 0.19 ± 0.03 | -0.47 | ns | 1 | 0.64 |
| v. saturation (%) | 16 | 20 | 0.019 ± 0.003 | 0.023 ± 0.006 | 2.38 | ns | 0.14 | 0.02 |

**LAB**

| trait | N (forest) | N (prairie) | forest (mean ± s.d.) | prairie (mean ± s.d.) | t statistic | | p (adjusted) | p (initial) |
|---|---|---|---|---|---|---|---|---|
| body (mm) | 20 | 20 | 88.2 ± 3.5 | 84.8 ± 3.1 | -3.25 | * | 0.0148 | 0.0025 |
| weight (g) | 20 | 20 | 1.32 ± 0.1 | 1.3 ± 0.08 | -0.72 | ns | 0.75 | 0.48 |
| ear (mm) | 20 | 20 | 18.0 ± 0.6 | 16.9 ± 1.2 | -3.28 | * | 0.0148 | 0.0028 |
| tail (mm) | 20 | 20 | 106.8 ± 6.4 | 73.3 ± 5.0 | -18.42 | *** | 1.17E-18 | 8.37E-20 |
| hindfoot (mm) | 20 | 20 | 23.2 ± 0.9 | 20.8 ± 0.9 | -8.32 | *** | 3.92E-09 | 4.36E-10 |
| d. brightness (a.u.) | 31 | 31 | 24.1 ± 6.3 | 43.0 ± 7.0 | 11.16 | *** | 4.54E-15 | 3.49E-16 |
| f. brightness (a.u.) | 31 | 31 | 53.2 ± 9.2 | 82.0 ± 12.2 | 10.5 | *** | 9.81E-14 | 8.17E-15 |
| v. brightness (a.u.) | 31 | 31 | 134.2 ± 25.7 | 191.6 ± 26.4 | 8.68 | *** | 3.71E-11 | 3.38E-12 |
| d. hue (degrees) | 31 | 31 | 37.8 ± 2.4 | 42.7 ± 2.4 | 8.03 | *** | 4.38E-10 | 4.38E-11 |
| f. hue (degrees) | 31 | 31 | 41.9 ± 1.8 | 45.6 ± 2.5 | 6.77 | *** | 6.98E-08 | 9.97E-09 |
| v. hue (degrees) | 31 | 31 | 58.6 ± 4.3 | 65.7 ± 3.6 | 6.97 | *** | 2.62E-08 | 3.28E-09 |
| d. saturation (%) | 31 | 31 | 0.23 ± 0.05 | 0.25 ± 0.03 | 1.86 | ns | 0.27 | 0.07 |
| f. saturation (%) | 31 | 31 | 0.22 ± 0.03 | 0.21 ± 0.04 | -0.89 | ns | 0.75 | 0.38 |
| v. saturation (%) | 31 | 31 | 0.021 ± 0.004 | 0.022 ± 0.004 | 1.68 | ns | 0.3 | 0.1 |

**Table S2.**

**QTL effect sizes.** Model results using fitqtl for each trait. Abbreviations: chr. = chromosome; LOD = log of the odds score; pos. = position in basepairs (bp); CI = 95% Bayes' credible interval (bounds given in bp, width given in Mb); a = additive effect (estimate ± standard error); d = dominance effect (estimate ± standard error); abs = absolute value; PVE = percent variance explained; * = QTL is transgressive; pgm = cross direction. For tail models, effect sizes were estimated while including all significant loci (left) or in models with each locus alone (right). Note that linear models strongly supported the presence of all five loci ($F_{2,540}$ comparing a full model with all 5 loci vs models with one locus dropped: 10.6 - 49.0). Labels indicate trait of interest, with any additive covariates in parentheses.

| **Primary models** | Trait | **Tail (body)**<br>% variance explained (all significant QTL): 27.3% | | | | |
|---|---|---|---|---|---|---|
| | chr. | 6 | 7 | 12 | 14 | 15 |
| | LOD | 4.9 | 4.7 | 5.3 | 5.1 | 17.4 |
| | pos. | 97457639 | 37729627 | 17493789 | 229987 | 41332882 |
| | CI | (77197532 , 120898287) | (22930804 , 118666602) | (10643762 , 24185533) | (220736 , 41048679) | (406222 , 41452107) |
| | CI width | 43.7 | 95.7 | 13.5 | 40.8 | 41 |
| Models including all significant QTL // one QTL alone | a | 1.17 ± 0.26 // 1.41 ± 0.3 | 1.28 ± 0.26 // 1.37 ± 0.3 | 1.6 ± 0.27 // 1.52 ± 0.31 | 1.59 ± 0.28 // 1.55 ± 0.32 | 2.71 ± 0.27 // 2.76 ± 0.3 |
| | d | -0.29 ± 0.38 // -0.2 ± 0.43 | 0.18 ± 0.38 // 0.33 ± 0.44 | 0.19 ± 0.38 // 0.20 ± 0.44 | 0.14 ± 0.38 // 0.37 ± 0.44 | 0.50 ± 0.38 // 0.50 ± 0.41 |
| | abs. d/a | 0.25 // 0.14 | 0.14 // 0.24 | 0.12 // 0.13 | 0.09 // 0.24 | 0.19 // 0.18 |
| | PVE | 2.6 // 3.8 | 3 // 3.6 | 4.5 // 4.1 | 4 // 3.9 | 12.1 // 12.8 |
| **Models with additional covariates** | Trait | **Tail (body, sex, age)** | | | | |
| | chr. | 6 | 7 | 12 | 14 | 15 |
| | LOD | 5.4 | 5 | 6.2 | 4.3 | 18.7 |
| | pos. | 97457639 | 37681732 | 16589347 | 229987 | 41332882 |
| | CI | (77183524 , 113132464) | (23423646 , 86976095) | (11903424 , 22687967) | (220736 , 41073607) | (473899 , 41465979) |
| | CI width | 35.9 | 63.6 | 10.8 | 40.9 | 41 |
| Models including all significant QTL // one QTL alone | a | 1.24 ± 0.26 // 1.46 ± 0.3 | 1.31 ± 0.25 // 1.39 ± 0.29 | 1.74 ± 0.27 // 1.66 ± 0.31 | 1.37 ± 0.28 // 1.40 ± 0.33 | 2.76 ± 0.27 // 2.81 ± 0.29 |
| | d | -0.30 ± 0.37 // -0.23 ± 0.43 | 0.28 ± 0.37 // 0.43 ± 0.43 | 0.02 ± 0.37 // -0.06 ± 0.43 | 0.26 ± 0.37 // 0.46 ± 0.44 | 0.48 ± 0.37 // 0.47 ± 0.4 |
| | abs. d/a | 0.24 // 0.16 | 0.21 // 0.31 | 0.01 // 0.04 | 0.19 // 0.33 | 0.18 // 0.17 |
| | PVE | 2.9 // 4.1 | 3.2 // 3.8 | 5.1 // 4.7 | 2.9 // 3.2 | 12.5 // 13.2 |

| **Primary models** | Trait | **Dorsal hue** | **Flank hue** | **Flank brightness** |
|---|---|---|---|---|
| | chr. | 15 | 15 | 21* |
| | LOD | 60.1 | 71.5 | 4 |
| | pos. | 11545604 | 21979520 | 35087484 |
| | CI | (406222 , 40539040) | (406222 , 39415731) | (25040451 , 69570542) |
| | CI width | 40.1 | 39 | 44.5 |
| | a | -1.96 ± 0.11 | -1.93 ± 0.09 | 3.47 ± 1.07 |
| | d | 0.52 ± 0.15 | 0.58 ± 0.13 | -4.32 ± 1.51 |
| | abs. d/a | 0.26 | 0.3 | 1.25 |
| | PVE | 40.0 | 45.6 | 3.4 |
| **Models with additional covariates** | Trait | **Dorsal hue (pgm)** | **Flank hue (pgm)** | **Flank brightness (pgm)** |
| | chr. | 15 | 15 | 21* |
| | LOD | 67.2 | 78.1 | 4.2 |
| | pos. | 11545604 | 10474970 | 35087484 |
| | CI | (403562 , 40734548) | (475455 , 40590260) | (27589255 , 69570542) |
| | CI width | 40.3 | 40.1 | 42 |
| | a | -1.95 ± 0.1 | -1.92 ± 0.09 | 3.17 ± 1.04 |
| | d | 0.54 ± 0.14 | 0.59 ± 0.12 | -4.73 ± 1.47 |
| | abs. d/a | 0.28 | 0.31 | 1.49 |
| | PVE | 39.7 | 45.3 | 3.3 |

### Table S3.

**Taqman assay probes.** Custom Taqman SNP genotyping assays used to genotype four SNPs differentiating the inversion allele from the reference allele.

| Taqman assay name | SNP location | Forward primer | Reverse primer | Probe | Reference allele | Inversion allele |
|---|---|---|---|---|---|---|
| March6 | chr15_8729600 | AGCCCACCAAGCCATACTC | AAGGCCGGCGGTAAGG | TGAAAGCC[A/G]ACAGGTC | A - VIC | G - FAM |
| Zfp622 | chr15_14131681 | TCCATGGTTTCATCATCACATTCCA | GCAGGACCCTAGTGAGATTGAG | TTTTCGGA[G/A]AATAAC | G - VIC | A - FAM |
| Slc45a2 | chr15_23939629 | TGCAGCAGGAATCCCAAGATG | GCCTGCCCAAGAGTTTGTACA | CATGGTGTGG[C/T]TCCTAA | C - VIC | T - FAM |
| Skp2 | chr15_29639255 | CTGTCTGAGTGCTCCAAGCT | ACAATGGGATCCGAAAGTTGCA | CTTCCAG[G/A]CTTAGATTC | G - VIC | A - FAM |

**Table S4.**

**Groups used to classify habitat type.** Habitat Group = grouping used for analysis, after binning categories across age groups. Habitat = habitat names used in the source model. Value = numeric code used in the source model.

| Habitat Group | Habitat | Value |
|---|---|---|
| alpine | Alpine | 20 |
| | Subalpine Parkland | 35 |
| aspen | Quaking Aspen | 23 |
| cliffs_canyons | Cliffs and Canyons | 62 |
| coastal_spruce_cedar_redwood | Coastal Spruce, Cedar or Redwood mature | 63 |
| | Coastal Spruce, Cedar or Redwood medium | 66 |
| | Coastal Spruce, Cedar or Redwood old-growth | 72 |
| | Coastal Spruce, Cedar or Redwood young | 73 |
| crops_and_developed | Urban (High Intensity Developed) | 32 |
| | Suburban (Moderate Intensity Developed) | 56 |
| | Cultivated Crops | 57 |
| | Rural Residential (Low Intensity Developed) | 61 |
| douglas_fir_western_hemlock | Douglas Fir - Western Hemlock mature | 14 |
| | Douglas Fir - Western Hemlock medium | 33 |
| | Douglas Fir - Western Hemlock young | 59 |
| | Douglas Fir - Western Hemlock old-growth | 75 |
| grassland | Montane Grasslands and Dry Meadows | 16 |
| | Alkali and Desert Grasslands | 26 |
| | Coastal and Valley Grasslands | 36 |
| | Pasture or Hay | 58 |
| | Exotic Grasslands and Annuals | 65 |
| | Columbia Basin Grasslands and Prairie | 74 |
| juniper | Western Juniper | 39 |
| lava_dunes_playa_burns_beaches | Coastal Dunes and Beaches | 24 |
| | Inland Dunes | 25 |
| | Lava | 29 |
| | Playa and Barren Ash | 30 |
| | Rocky Coast | 31 |
| | Burns | 69 |
| lodgepole_pine | Lodgepole Pine mature | 5 |
| | Lodgepole Pine young | 45 |
| mixed_conifer | Mixed Conifer (White or Douglas Fir/Pine) mature | 3 |
| | Mixed Conifer (White or Douglas Fir/Pine) medium | 4 |
| | Mixed Conifer (White or Douglas Fir/Pine) old-growth | 21 |
| | Mixed Conifer (White or Douglas Fir/Pine) young | 40 |
| mixed_hardwood_conifer | Mixed Hardwood - Conifer mature | 12 |
| | Mixed Hardwood - Conifer old-growth | 13 |
| | Mixed Hardwood - Conifer medium | 18 |
| | Mixed Hardwood - Conifer young | 44 |
| oak | Oak | 34 |
| | Mixed Oak - Conifer mature | 41 |
| | Mixed Oak - Conifer old-growth | 42 |
| | Mixed Oak - Conifer young to medium | 43 |
| ponderosa | Ponderosa Pine medium | 8 |
| | Ponderosa Pine mature | 46 |
| | Ponderosa Pine old-growth | 47 |
| | Ponderosa Pine young | 48 |
| sagebrush | Big Sagebrush fair - good | 2 |
| | Low Sagebrush fair - good | 9 |
| | Mountain Big Sagebrush fair - good | 10 |
| | Big Sagebrush poor | 71 |
| | Low Sagebrush poor | 76 |
| | Mountain Big Sagebrush poor | 77 |
| shrub | Salt Desert Scrub | 11 |
| | Early Shrub-Tree | 17 |
| | Chaparral | 19 |
| | Canyon & Montane Shrubland | 22 |
| silver_fir_mtn_hemlock | Silver Fir - Mountain Hemlock medium | 6 |
| | Silver Fir - Mountain Hemlock mature | 49 |
| | Silver Fir - Mountain Hemlock old-growth | 50 |
| | Silver Fir - Mountain Hemlock young | 51 |
| siskiyou | Siskiyou Mixed Conifer medium | 7 |
| | Siskiyou Mixed Conifer old-growth | 52 |
| | Siskiyou Mixed Conifer mature | 53 |
| | Siskiyou Mixed Conifer young | 54 |
| spruce | Spruce - Subalpine Fir old-growth | 1 |
| | Spruce - Subalpine Fir young | 55 |
| | Spruce - Subalpine Fir medium to mature | 64 |
| water_riparian | Marshes, Bogs and Emergent Wetlands | 15 |
| | Open Water (Big Rivers and Reservoirs) | 27 |
| | Saltmarsh | 28 |
| | Coastal and Valley Riparian | 37 |
| | Montane Wetlands | 38 |
| | Interior Lowland and Foothill Riparian | 67 |
| | Lowland Woody Wetlands and Swamps | 68 |
| | Bays and Estuaries | 70 |

**Data S1. (separate file)**

**Wild-caught specimens from this study.** Names (internal to this study; Location and Site) and locations (WGS84, Latitude and Longitude) of trapping sites. Elevation_m: elevation of each site in meters. Transect_distance_km: east-west distance along the transect in kilometers east of the central site. Harvard_MCZ_accession: accession number of each specimen at the Harvard University Museum of Comparative Zoology. Study_ID: study-specific ID number. Age: age class (A = adult, S = subadult, J = juvenile). Sex: M = male, F = female. * = used as colony founder, ** = used for whole genome sequencing.

**Data S2. (separate file)**

**Peromyscus phylogeny sample ids and museum collections.** Study-specific ID (id) with corresponding species/subspecies (species) of samples included in the *Peromyscus* phylogenies. Samples were from different museum collections (collection), with corresponding museum IDs (museum_id) listed.

**Data S3. (separate file)**

**Genotypes and phenotypes for mice from across North America.** Study-specific ID (id) for mice used to determine the distribution of the inversion across North America. SRA.Run: NCBI SRA run number for samples obtained from published datasets. Other_ID: ID used in original publication or museum databases. Museum.accession: museum ID if applicable (MCZ = Harvard University Museum of Comparative Zoology, MVZ = University of California Berkeley Museum of Vertebrate Zoology, UWBM = University of Washington Burke Museum). Longitude, Latitude: coordinates for location of capture site. Dataset: origin of sequencing data or tissue. Sequencing.type: sequencing approach used for published datasets, focal mice or museum samples. Inversion_genotype: inferred genotype at the inversion (ref/ref = homozygous reference allele, inv/inv = homozygous inversion allele). Percent_forested: percent of pixels in 1-km radius around capture site categorized as forest habitat. Standard morphological measurements recorded for body length (body_length) and tail length (tail_length). State/region: region of capture site. Population_number: population number corresponding to Figure 5F,G.

**Data S4. (separate file)**

**Wild-caught forest, prairie and transect mice phenotypes**. Study-specific ID number (id, field_id), trap site name (location, site), location (latitude, longitude), and elevation (in feet, elevation.ft, and meters, elevation.m) of wild-caught forest and prairie and Cascades transect mice (HUMCZ_ID = Harvard University Museum of Comparative Zoology accession number), with phenotypes. Also included are the transect specimens from the University of Washington's Burke Museum. Sex: M = male, F = female. Age: age class, A = adult, S = subadult, J = juvenile. Pregnant: whether the mouse was (Y) or was not (N) visibly pregnant at capture; we excluded pregnant individuals when analyzing body weight. Coat color phenotypes: brightness, saturation, and hue taken as the median of 3-5 measurements from each body region (D = dorsal stripe, F = flank, V = ventrum). Body measurements: tail, body ( = total – tail), ear, and hindfoot lengths in millimeters and weight in grams. Weight_log10 = $\log_{10}$-transformed weights used for analysis; tail.to.body = tail length / body length. Mice used as colony founders are identified (colony_founder = TRUE), and subspecies (P. m. gambelii or P. m. rubidus), ecotype (prairie or forest) and transect position (dist_east, in kilometers east of the central site) are provided. For mice with sequencing data, ngsadmix_forest_ancestry gives genome-wide forest ancestry determined by ngsAdmix and inversion_genotype denotes whether mice were determined to be homozygous for the inversion (inv/inv), heterozygous (inv/ref) or homozygous for the reference (ref/ref) allele.

**Data S5. (separate file)**

**Lab-born forest and prairie body dimensions.** Laboratory ID (id), subspecies and ecotype (*P. m. gambelii* = prairie, *P. m. rubidus* = forest), sex (M = male, F = female), and age (in days) of laboratory-born mice, as well as body dimensions (tail length, body length = total – tail, ear, and hindfoot length in millimeters, weight in grams, and $\log_{10}$-transformed weight values).

**Data S6. (separate file)**

**Lab-born forest and prairie coat color phenotypes.** Laboratory ID (id), subspecies and ecotype (*P. m. gambelii* = prairie, *P. m. rubidus* = forest), and coat color phenotypes of 60-70 day old laboratory-born mice. Coat color phenotypes: brightness, saturation, and hue taken as the median of 3-5 measurements from each body region (D = dorsal stripe, F = flank, V = ventrum).

**Data S7. (separate file)**

**F2 hybrid phenotypes.** Study ID (ID, numeric, and id, character), sex (F = female, M = male; see also sex.male column), birth date (DOB = MM/DD/YY), age in days, and phenotype information for second-generation forest-prairie intercross hybrids (type = F2). Tail = tail length in millimeters (mm), total = total length (nose to tail tip) in mm, weight = weight in grams, foot.std = hindfoot length in mm, body = body length (total – tail) in mm. Transformed values: weight_log10 = $\log_{10}$-transformed weights; tail.resid.body = residual of tail length from a linear model *tail ~ body* using all F2s; foot.std.resid.body = residual of foot length from a linear model *foot.std ~ body* using all F2s. Coat color phenotypes: brightness, saturation, and hue taken as the median of 3-5 measurements from each body region (D = dorsal stripe, F = flank, V = ventrum). Some genetic information is also included: pgm = cross direction (0 = forest female x prairie male, 1 = prairie female x forest male); family = family identifier, mice with the same value are full siblings. Total_forest (total_prairie, total_het) = percent of genome determined to be homozygous for forest ancestry (prairie ancestry, heterozygous ancestry). Total_forest_no15 (total_prairie_no15, total_het_no15) provides the same information excluding all of chromosome 15. Total_forest_alleles_no15 = percent of alleles genome-wide determined to be forest ancestry, excluding chromosome 15. Eight individuals with too low coverage sequencing data to be used for genetic mapping are marked with geno.too.low.coverage = Y.

**Data S8. (separate file)**

**Habitat category by site across transect.** Proportion of the area (proportion_area) within a radius (radius_km) of 0.5, 1, or 2 kilometers of each capture site (uniquely identified by the study-specific identifiers Location and Site, combined into one value in Full_Location) located at each position along the transect (distance_east = distance east of the central site in kilometers) identified as each habitat (Habitat = category directly from the Oregon Biodiversity Information Center models) and binned habitat group (defined as in Table S4). Region bins sites into related groups (0_west = forest, 1_mid = central Cascades transect, 1.5_new = additional museum specimens, 2_east = prairie).

**Data S9. (separate file)**

Soil color by site across transect. Proportion of the area (pct_color_in_site) within a radius (radius_km) of 0.5, 1, or 2 kilometers of each capture site (uniquely identified by the study-specific identifiers Location and Site) and assuming specific moisture conditions (condition = moist, dry, or the typical moisture at that site, see Materials and Methods) categorized into each Munsell-scale color (munsell). Munsell_hue = hue only; munsell_value = value only; munsell_chroma = chroma only; munsell_hue_num = munsell_hue converted to degrees as described in Materials and Methods.
